# Accounting for variability in ion current recordings using a mathematical model of artefacts in voltage-clamp experiments

**DOI:** 10.1101/2019.12.20.884353

**Authors:** Chon Lok Lei, Michael Clerx, Dominic G. Whittaker, David J. Gavaghan, Teun P. de Boer, Gary R. Mirams

## Abstract

Mathematical models of ion channels, which constitute indispensable components of action potential models, are commonly constructed by fitting to whole-cell patch-clamp data. In a previous study we fitted cell-specific models to hERG1a (Kv11.1) recordings simultaneously measured using an automated high-throughput system, and studied cell-cell variability by inspecting the resulting model parameters. However, the origin of the observed variability was not identified. Here we study the source of variability by constructing a model that describes not just ion current dynamics, but the entire voltage-clamp experiment. The experimental artefact components of the model include: series resistance, membrane and pipette capacitance, voltage offsets, imperfect compensations made by the amplifier for these phenomena, and leak current. In this model, variability in the observations can be explained by either cell properties, measurement artefacts, or both. Remarkably, by assuming that variability arises exclusively from measurement artefacts, it is possible to explain a larger amount of the observed variability than when assuming cell-specific ion current kinetics. This assumption also leads to a smaller number of model parameters. This result suggests that most of the observed variability in patch-clamp data measured under the same conditions is caused by experimental artefacts, and hence can be compensated for in post-processing by using our model for the patch-clamp experiment. This study has implications for the question of the extent to which cell-cell variability in ion channel kinetics exists, and opens up routes for better correction of artefacts in patch-clamp data.

## 1 Introduction

Mathematical modelling and computational simulations have been remarkably successful in providing mechanistic insight into many electrophysiological phenomena. Quantitative models of the action potential have demonstrated their usefulness in basic research and are beginning to be used in safety-critical applications Mirams et al. (2012); Niederer et al. (2018); Li et al. (2019). Mathematical models of ion channels constitute indispensable components of these action potential models. Even when models are fitted to the best available data, uncertainty in their parameter values remains, which can be due to measurement uncertainty and/or physiological variability Mirams et al. (2016). Thus, identifying and quantifying sources of uncertainty enables informed decision making when using models in safety-critical applications US National Research Council (2012).

Whole-cell patch-clamp experiments (in voltage-clamp configuration) are a common source of data for calibrating ion channel models. To study the dynamics of ion channels, currents through the cell membrane are often measured with a patch-clamp amplifier. In voltage-clamp mode, a patch-clamp amplifier is a sensitive feedback amplifier that rapidly calculates, applies and reports the small currents necessary to maintain a given voltage across a cell’s membrane (and vice versa for current-clamp mode) Sigworth (1995b). Typical peak current magnitudes are on the order of pA to μA, depending on the size and type of cell; the voltage across the cell membrane (potential inside minus outside) typically is within the range −140 to +60 mV.

We use the term ‘*kinetics*’ to describe the voltage-dependent opening and closing of ion channels in response to changes in membrane voltage. ‘*Maximal conductance*’ determines the magnitude of the current that would flow if all the channels were open, a value proportional to the number of channels in the membrane. High levels of variability in kinetics have been observed in many studies Bekkers et al. (1990); Finkel et al. (2006); Feigenspan et al. (2010); Golowasch (2014); Santillo et al. (2014); Altomare et al. (2015); Annecchino and Schultz (2018). A previous study Lei et al. (2019b) analysed hERG current kinetics recorded simultaneously in 124 cells using an automated high-throughput patch-clamp machine. The experiments used Chinese hamster ovary (CHO) cells stably expressing hERG1a. Since the cells expressed the same genes and were measured at the same time under highly similar conditions, one might expect the resulting current kinetics to be very similar across different cells. However, a high level of variability was seen, similar to that in previous studies using manual patch-clamp experiments conducted one cell at a time over several days Beattie et al. (2018). This raises a question — what is the origin of the observed variability?

Figure 1 (left) shows an idealised voltage-clamp experiment, where the cell is connected directly to an ammeter which records the current of interest, *I*_ion_, while clamping the membrane to the command voltage, *V*_cmd_ (its equivalent circuit is shown Supplementary Figure S1). In other words, a perfect patch-clamp experiment can be represented with the following simple equations/assumptions:

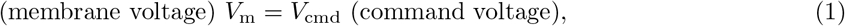

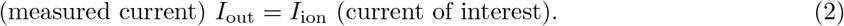

**Figure 1:**
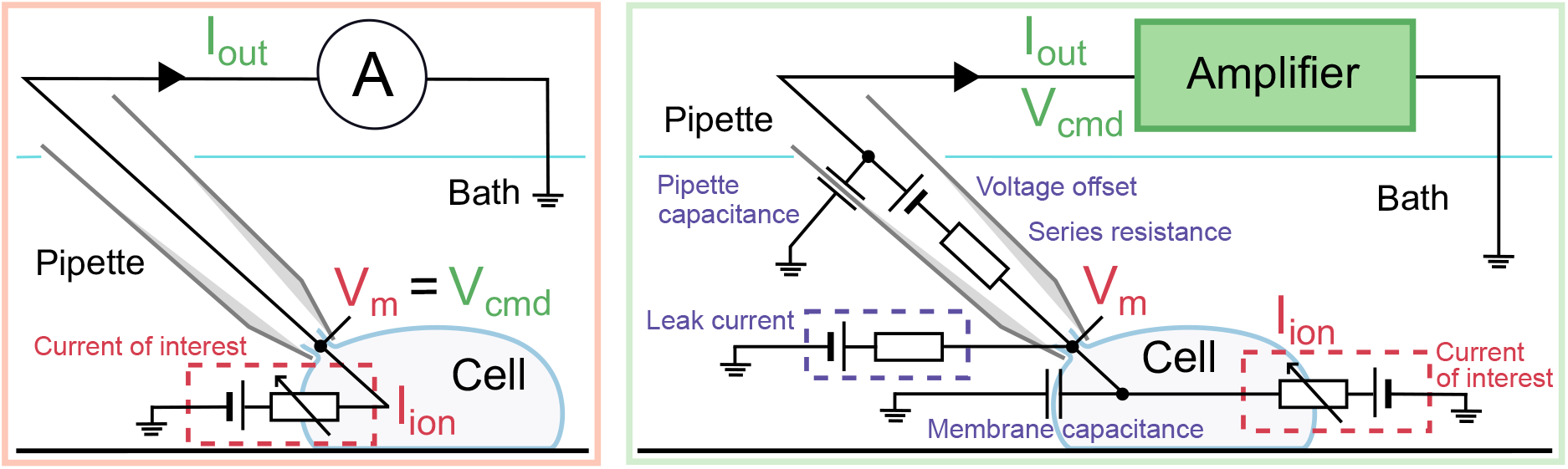
Schematics of the voltage-clamp experiment: **(left)** idealised and **(right)** more realistic. **Left:** Only the current of interest (red) is modelled. The membrane voltage, *V*_m_ (red) is assumed to be the same as the command voltage, *V*_cmd_ (green) set by the amplifier, and the observed current, *I*_out_ (green) is assumed to be equal to the ion channel current, *I*_ion_ (red). **Right:** Here, not only the current of interest (red) is modelled, but the patch-clamp amplifier process (green) and all of the experimental artefacts (purple) are included in the model. The differences between *V*_m_ (red) and *V*_cmd_ (green), and between *I*_ion_ (red) and *I*_out_ (green) are explicitly modelled in this framework.

This is the first of three ways of modelling the patch-clamp system that will be discussed in this manuscript (each enclosed in boxes as above).

In realistic voltage-clamp experiments, as illustrated in Figure 1 (right), the cell membrane acts as a capacitor in parallel to the ion currents. Between the pipette electrode and the cell, there is a *series resistance*, which induces a mismatch between the membrane voltage *V*_m_ and the command voltage *V*_cmd_, causing a shift in measured current-voltage relationships Marty and Neher (1995); Sherman et al. (1999). Furthermore, there is a voltage offset introduced between the pipette electrode and the cell Neher (1995); and the wall of the pipette (or the well plate in an automatic system) behaves as a capacitor. Finally, a finite seal resistance can cause a substantial leak current that contaminates the recording of the current of interest. All of these can be *partially* compensated by the patch-clamp amplifier using real-time hardware adjustments, or addressed in post-processing, and the remainder are what we term ‘*experimental artefacts*’. Although real-world experiments suffer from these remaining artefacts many studies assume the experimental apparatus perfectly compensates for any discrepancies, so the idealised assumptions (Eqs. (1)–(2)) are frequently used when analysing experimental data. Often the existence of a leak current is acknowledged and an estimate for it is subtracted in a post-processing step to modify the *I*_out_ in Eq. (2) as we will discuss later.

A previous microelectrode study Raba et al. (2013) estimated and subtracted a transient cell membrane capacitive current post-hoc using a mathematical model that included many of these components. In this paper, we relax further the typical, ideal voltage-clamp assumptions by introducing a mathematical model that allows and accounts for artefacts, imperfect amplifier compensations for artefacts, together with any residual uncompensated leak current. We then validate the mathematical model experimentally using electrical model cells, for which we designed a new type of electrical model cell which exhibits simple dynamic behaviour. Using this new mathematical model, variability in observations can be attributed to measurement artefacts as well as current properties. We develop an algorithm to optimise the ion current maximal conductances, a set of current kinetic parameters and measurement artefact parameters for more than one cell at the same time. Finally, we compare fits and assess the predictions of models calibrated with either ideal voltage-clamp assumptions or realistic voltage-clamp assumptions. Whilst different cells have varying maximal current conductance, the study has implications for the significance or even existence of cell-cell variability in ion channel kinetics, and opens up routes for better correction of artefacts in patch-clamp data.

## 2 A detailed mathematical model of a voltage-clamp experiment

Figure 2 presents an expanded circuit for the more realistic voltage-clamp experiment shown in Figure 1 (right), including the amplifier components that compensate for the additional artefacts in the realistic experiment. Our goal is to observe the ion current across the cell membrane, *I*_ion_. This current is present in the ‘Cell Model’ in Figure 2 (shown with black, dashed box). Between the Cell Model and the Headstage (green, dashed box) is where the pipette (or the well plate in an automatic system) sits, separating the cell membrane and the pipette electrode, which includes many of the undesired artefact components shown in Figure 1 (right). There are five main undesired effects in this voltage-clamp set-up: 1. parasitic/pipette capacitance (a capacitance effect induced by the pipette wall), 2. (cell) membrane capacitance, 3. series resistance (a lumped term for all resistances between the pipette electrode and the cell), 4. voltage offset (due to amplifier offsets, electrode offsets, junction potentials, etc.), 5. leak current (a current leak through the pipette-cell seal and any leak through the membrane). Table 1 contains a glossary of symbols and parameters used for these quantities throughout this paper.

**Table 1:** Glossary of symbols and parameters. We also denote a (machine or post-processing) estimate of a parameter *X* as *X*\*, and the error in the estimate of the same parameter as *X*^†^. The range of typical values are taken from Weerakoon et al. (2009); Neher (1995); HEKA Elektronik Dr. Schulze GmbH (2007–2018); Axon Instruments Inc. (1997–1999), unless otherwise specified.

| Symbol | Typical values | Description |
| --- | --- | --- |
| **General** |  |  |
| $t$ | — | Time variable |
| **Capacitance** |  |  |
| $C_f$ | 0.3 pF | Feedback shunt capacitance |
| $C_m$ | 5–20 pF | Membrane capacitance. These are typical values for the CHO cells used in this study Lei et al. (2019b); 100–400 pF has been measured for individual cardiomyocytes in various species Satoh et al. (1996) ($\sim 190$ pF for human Sakakibara et al. (1993)); and can be as high as 100 nF for large cells such as *Xenopus* oocytes Schmitt and Koepsell (2002). |
| $C_p$ | 3–5 pF | Parasitic capacitance at the pipette/well plate; also known as ‘pipette capacitance’ in manual patch |
| **Current** |  |  |
| $I_{in}$ | — | Voltage-clamp current |
| $I_{inj}$ | — | Injection current |
| $I_{ion}$ | $\mathcal{O}(1)$ nA | Whole-cell ion channel current (can reach $\mathcal{O}(1)$ μA for *Xenopus* oocytes Schmitt and Koepsell (2002)) |
| $I_{leak}$ | — | Leakage current through imperfect seal |
| $I_m$ | — | Membrane current |
| $I_{out}$ | — | Recorded current |
| $I_p$ | — | Current drawn by parasitic capacitance of the pipette/well plate |
| **Resistance** |  |  |
| $R_f$ | 25 MΩ | Feedback resistance |
| $R_s$ | 5–20 MΩ | Series resistance between the pipette electrode and the cell |
| $R_{seal}$ | 0.5–10 GΩ | Seal resistance of the pipette tip |
| **Time constant** |  |  |
| $\tau_a$ | 50–2000 μs | Membrane access time constant, $\tau_a = R_s C_m$ |
| $\tau_{sum}$ | 5–70 μs | Response time of the summing amplifier |
| $\tau_{clamp}$ | 0.8 μs | Voltage-clamp time constant |
| $\tau_z$ | 7.5 μs | Transconductor time constant, $\tau_z = R_f C_f$ |
| **Voltage** |  |  |
| $V_{cmd}$ | $\mathcal{O}(100)$ mV | Command voltage; follows the voltage-clamp protocols |
| $V_{clamp}$ | $\mathcal{O}(100)$ mV | Clamp voltage |
| $V_m$ | $\mathcal{O}(100)$ mV | Membrane potential |
| $V_{off}$ | $\mathcal{O}(10)$ mV | Offset voltage, such as amplifier offsets, electrode offsets, junction potentials, etc. |
| $V_{out}$ | $\mathcal{O}(100)$ mV | Recorded voltage, $V_{out} = I_{out} R_f$ |
| $V_p$ | $\mathcal{O}(100)$ mV | Pipette potential |

**Figure 2:**
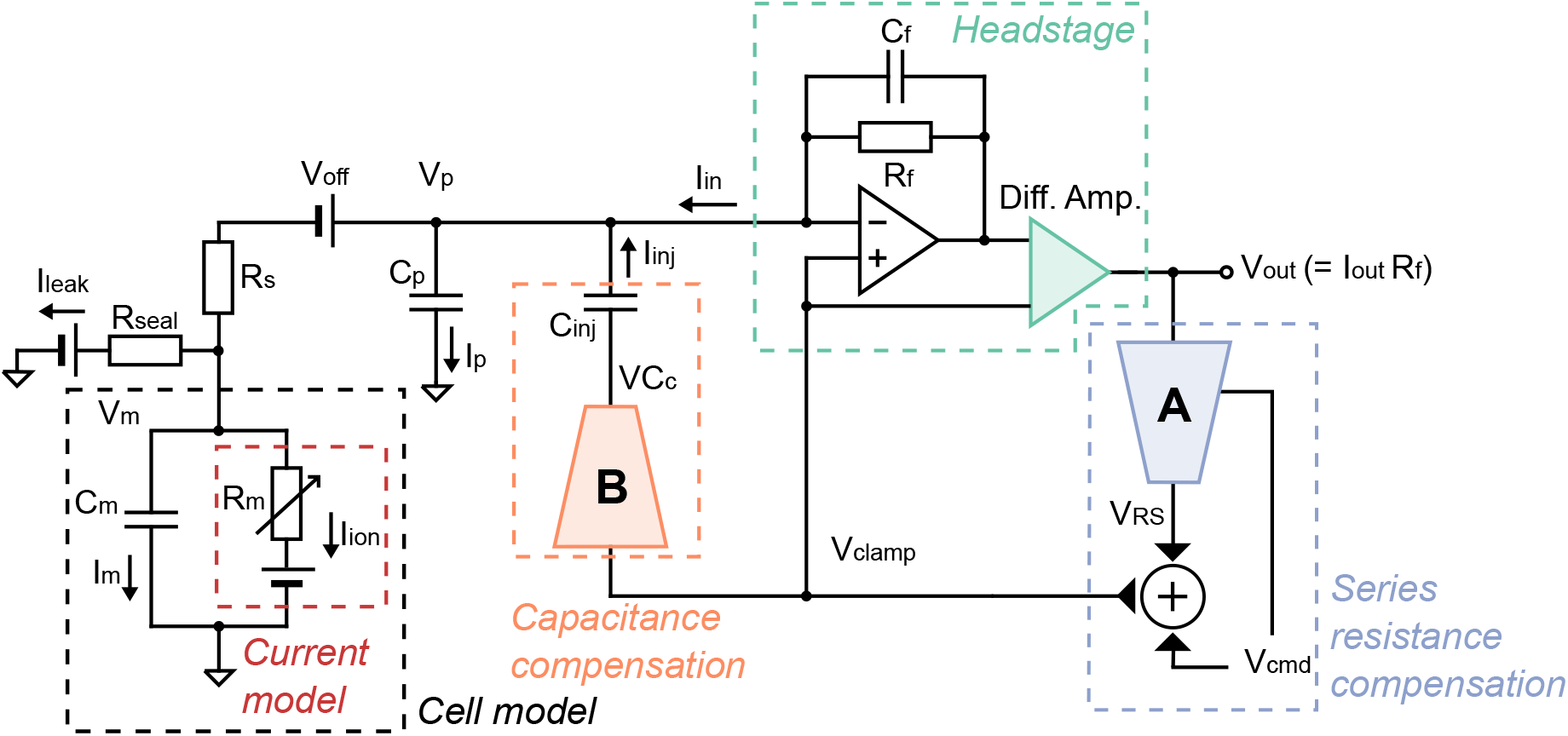
A more realistic voltage-clamp experiment equivalent circuit. This includes undesired factors such as voltage offset (*V*_off_), series resistance (*R*_s_) between the pipette electrode and the cell, cell capacitance (*C*_m_), pipette capacitance (*C*_p_), and leakage current (*I*_leak_), which can introduce artefacts to the recordings. The circuit also includes the components within a typical amplifier that are designed to compensate the artefacts. The blue **(A)** and orange **(B)** components are two idealised multiplying digital-to-analogue converters that control the amount of compensation. We assume that these, and the transimpedance amplifier and differential amplifier (green), to be ideal electrical components. Please refer to Table 1 for a description of the symbols.

To model the dynamics of the current of interest, *I*_ion_, the assumptions behind Eqs. (1)–(2) can be removed to instead model the entire voltage-clamp experiment and amplifier compensations with the equations below. Their derivation is given in Supplementary Section S2, where each of the undesired effects and how they are typically compensated is modelled based on published circuitry Moore et al. (1984); Neher (1992, 1995); Sigworth (1995a); Sigworth et al. (1995); Strickholm (1995); Sherman et al. (1999); Weerakoon et al. (2009, 2010); HEKA Elektronik Dr. Schulze GmbH (2007–2018); Axon Instruments Inc. (1997–1999).

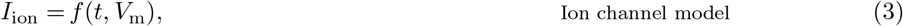

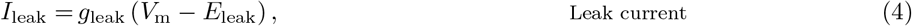

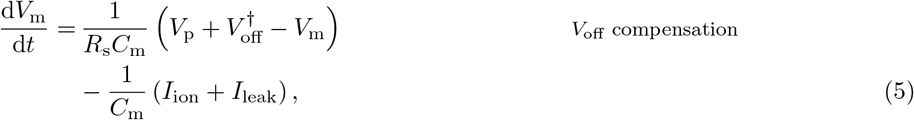

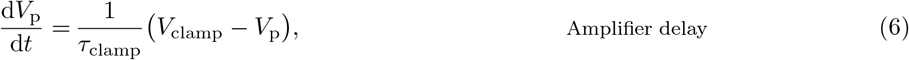

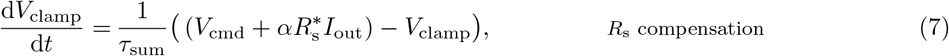

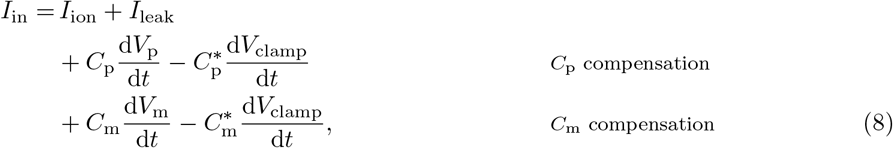

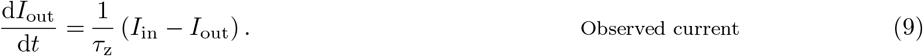

In these equations, *α* is the requested proportion of series resistance compensation (a machine setting, typically 70–85 %), and *g*_leak_ and *E*_leak_ are the conductance and the reversal potential of the leak current. The meaning of the remaining symbols is given in Table 1.

In the real cell data analysed later, a leak compensation was applied to the data itself, prior to model fitting. In whole-cell recordings, typically estimates (denoted by *) of the leak parameters in Eq. (4) are made in post-processing by assuming it is the only current active during a ‘leak step’ between voltages where *I*_ion_ ≈ 0. Finally, *I*_out_ is then adjusted by subtracting estimated leak

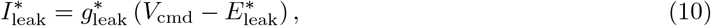

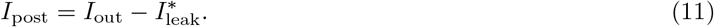

to give *I*_post_ as a better approximation of *I*_ion_.

## 3 Validating the mathematical model with electrical model cell experiments

Before applying the voltage-clamp experiment model to currents from a real biological cell, we test the performance of our mathematical model with *electrical model cells* — circuits made of electrical hardware components that mimic real cells. Some of these model cells are commercially available, e.g. HEKA MC 10, HEKA TESC, Axon or Molecular Devices PATCH-1U, and used to test and calibrate patch-clamp amplifiers.

We are interested in both the current readout of a voltage-clamp experiment and the membrane voltage that the cell experiences. Therefore we designed a circuit that connects the model cell to two amplifiers, one in voltage-clamp mode and one in current-clamp mode, as shown in Figure 3. This set-up can simultaneously perform the conventional voltage-clamp procedure on the model cell with one amplifier; whilst using the other in current-clamp mode (clamped to zero) to measure the clamped voltage at the terminal corresponding to the membrane via the current-clamp. Effectively, this set-up allows the membrane voltage *V*_m_ of the electrical model cell to be recorded whilst performing the conventional voltage-clamp current measurement.

**Figure 3:**
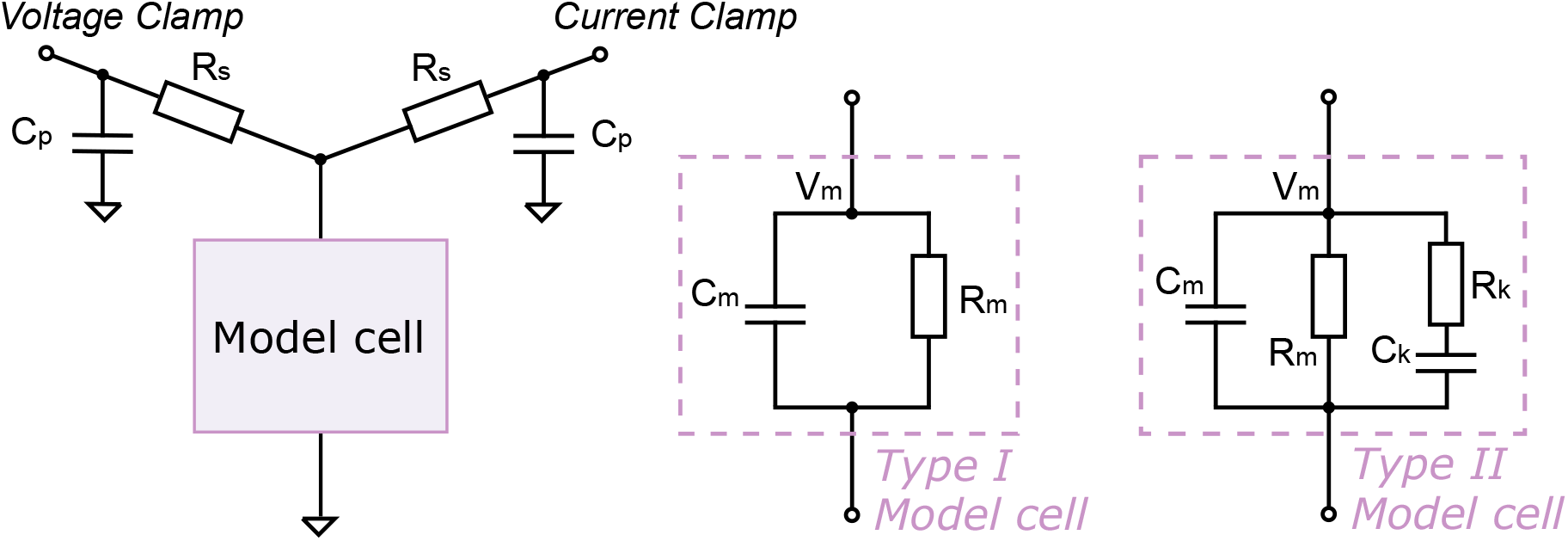
Circuit diagrams for electrical model cell experiments. **(Left)** A circuit set-up where a model cell is connected to both a voltage-clamp amplifier and a current-clamp amplifier. The voltage-clamp imposes a command voltage, *V*_cmd_, on the model cell and measures the current, *I*_out_; while the current-clamp simultaneously measures its ‘membrane voltage’, *V*_m_. **(Middle)** An equivalent circuit of the Type I Model Cell, which is identical to the commercial ‘black box’ model cells (HEKA TESC) under the ‘whole cell’ mode. **(Right)** An equivalent circuit for the Type II Model Cell. This model cell is designed to exhibit dynamics when stepping to different voltages, with a time constant similar to ionic currents. The circuits were built with discrete electrical components, with *C*_p_ = 4.7 pF, *R*_s_ = 30 MΩ, *C*_m_ = 22 pF, *R*_m_ = 500 MΩ, *C*_k_ = 1000 pF, and *R*_k_ = 100 MΩ.

### 3.1 Electrical model cell design

In Figure 3 (left and middle) is a circuit which is equivalent to commercially-available model cells (the design was based on the HEKA TESC), when under ‘whole cell’ mode. In this study, we call this a *Type I Model Cell*. It consists of a capacitor and a resistor in parallel to mimic the membrane capacitance, *C*_m_, and membrane resistance, *R*_m_. Unlike real ion channels, this simple electrical model cell lacks any current dynamics in the *R*_m_ resistor representing ion currents. We therefore developed a new type of model cell, termed a *Type II Model Cell*, which exhibits simple current dynamics when stepping to different voltages.

Figure 3 (right) shows the equivalent circuit of our Type II Model Cell. In addition to the usual *C*_m_ and *R*_m_ connected in parallel, this model cell has an extra component (*R*_k_ in series with *C*_k_) connected in parallel, to mimic the addition of another ion current with some kinetic properties. The time constant (*τ_k_* = *R*_k_*C*_k_) for this extra component was chosen to be 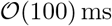, which is of the same order of magnitude as *I*_Kr_ dynamics. The dynamics of the Type II Model Cell allow us to test and understand the effects of series resistance, etc., and enable us to validate our mathematical voltage-clamp experiment model experimentally to check we have correctly modelled amplifier compensations.

### 3.2 Validation of the mathematical model

The experimental recordings using the simultaneous voltage clamp & current clamp set-up are shown in Figure 4 (solid lines), with Type I (A, C) and Type II (B, D) Model Cells. The measurements were performed with a holding potential at 0 mV.

**Figure 4:**
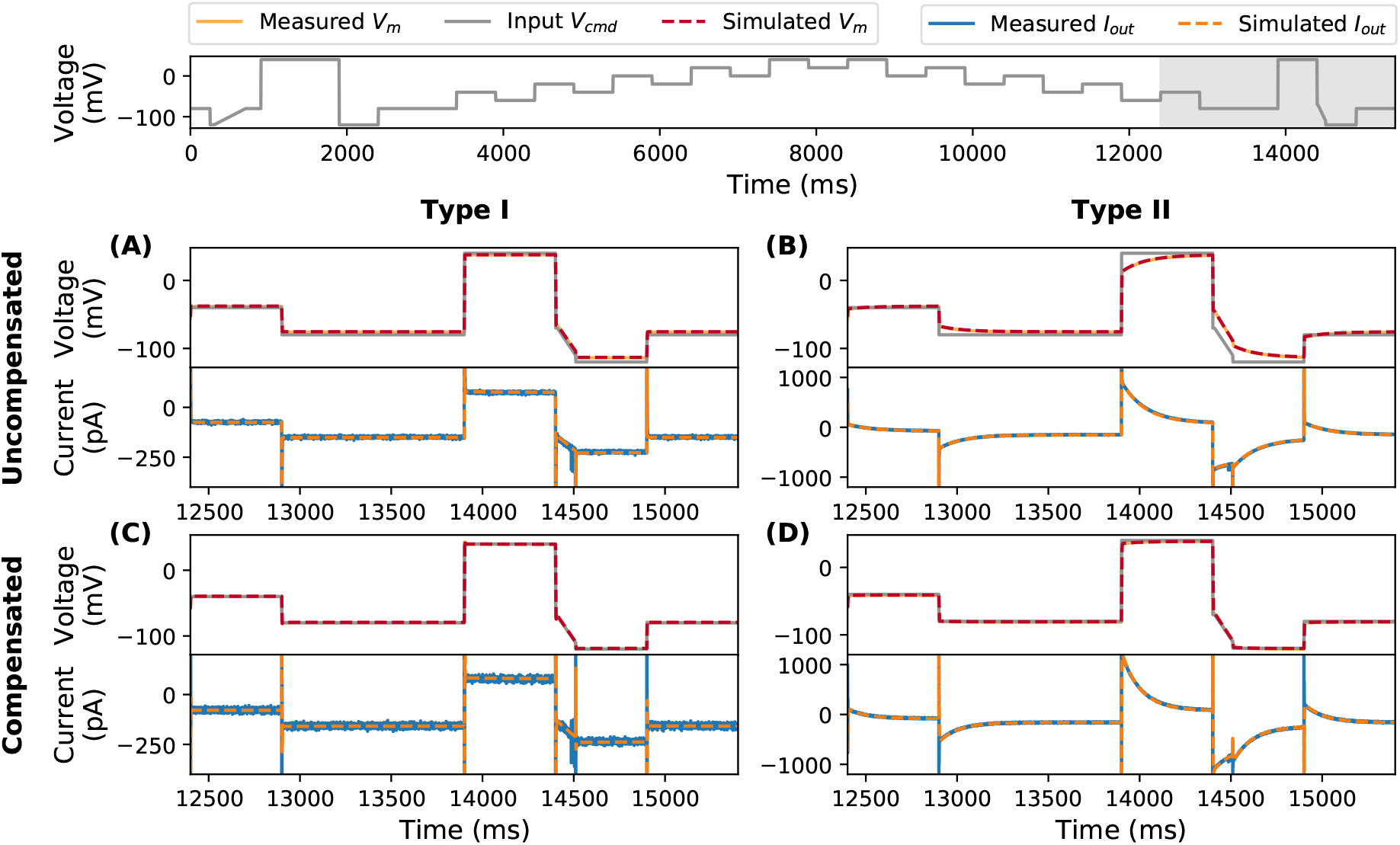
Model simulations (dashed lines) using the amplifier settings compared against the simultaneous voltage clamp-current clamp measurements of the model cells (solid lines). Measurements are shown without compensation using **(A)** Type I Model Cell and **(B)** Type II Model Cell; and measurements with automatic amplifier compensation for *V*_off_, *C*_p_, *C*_m_, and *R*_s_ with *α* = 80% using **(C)** Type I Model Cell and **(D)** Type II Model Cell. All command voltages were set to be the staircase voltage protocol Lei et al. (2019b) (top panel); here only the last 3 s of the measurement is shown, the whole trace is shown in Supplementary Figure S4. In the top panel of each subfigure, the grey lines represent the command voltage *V*_cmd_, and the orange/red lines represent the membrane voltage *V*_m_; the bottom panel shows the current readout via the voltage-clamp, *I*_out_.

We performed two sets of experiments, firstly with no amplifier compensation, and secondly with automatic amplifier compensation using a computer controlled amplifier (HEKA EPC 10 Double Plus) where amplifier settings could be set with high precision. Here, automatic adjustment of the compensation settings, including *V*_off_, *C*_p_, *C*_m_, and *R*_s_, was performed using the HEKA Patchmaster software which closely follows the compensation procedure operators perform by hand on many manual patch-clamp amplifiers (pages 80–84 of the HEKA manual HEKA Elektronik Dr. Schulze GmbH (2005–2016)).

For the simulations, parameters were set in Eqs. (3)–(9) to correspond to each set of experiments: for no amplifier compensation {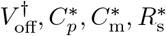, *g*_leak_} were set to zeros; while for the automatic amplifier compensation those parameters were set to the amplifier’s estimates. Results are shown in Figure 4 for part of the voltage clamp protocol to be used for *I*_Kr_ measurements later in this study (current under the full protocol is shown in Supplementary Figure S4).

Our model (dashed lines) is able to capture both the current and the membrane voltage very closely, for all these experiments. Note the differences between the membrane voltage *V*_m_ (grey) and the command voltage *V*_cmd_ (orange/red). For example, in the uncompensated case, due to the voltage drop across *R*_s_ in the Type I Model Cell *V*_m_ shows a simple offset. But in the Type II Model Cell *V*_m_ exhibits nonlinear dynamics while *V*_cmd_ does not. When the amplifier is actively compensating the differences between *V*_m_ and *V*_cmd_ were successfully reduced. All of these details are captured by the mathematical model so we are confident that Eqs. (3)–(9) are a good representation of the model cells, amplifier compensations and artefacts that occur when compensations are disabled or imperfect.

### 3.3 Parameter inference without compensations

Next, we attempt to use only the uncompensated, raw voltage-clamp measurements (i.e. only *I*_out_ in Figure 5A, and *V*_cmd_) to infer the underlying membrane voltage, *V*_m_, and the parameters of the model cells. We then compare the model *V*_m_ predictions with the current-clamp measurements. Here, focus is on the Type II Model Cell, since this should be the more challenging of the two and more similar to a real ionic current (similar results for the Type I Model Cell in Supplementary Section S5). To optimise the model parameters root-mean-squared error (RMSE) between the simulated and recorded *I*_out_ was minimised using a global optimisation algorithm Hansen (2006). All optimisation was done with an open source Python package, PINTS Clerx et al. (2019b), and simulations were performed in Myokit Clerx et al. (2016). All codes and data are freely available at https://github.com/CardiacModelling/VoltageClampModel.

**Figure 5:**
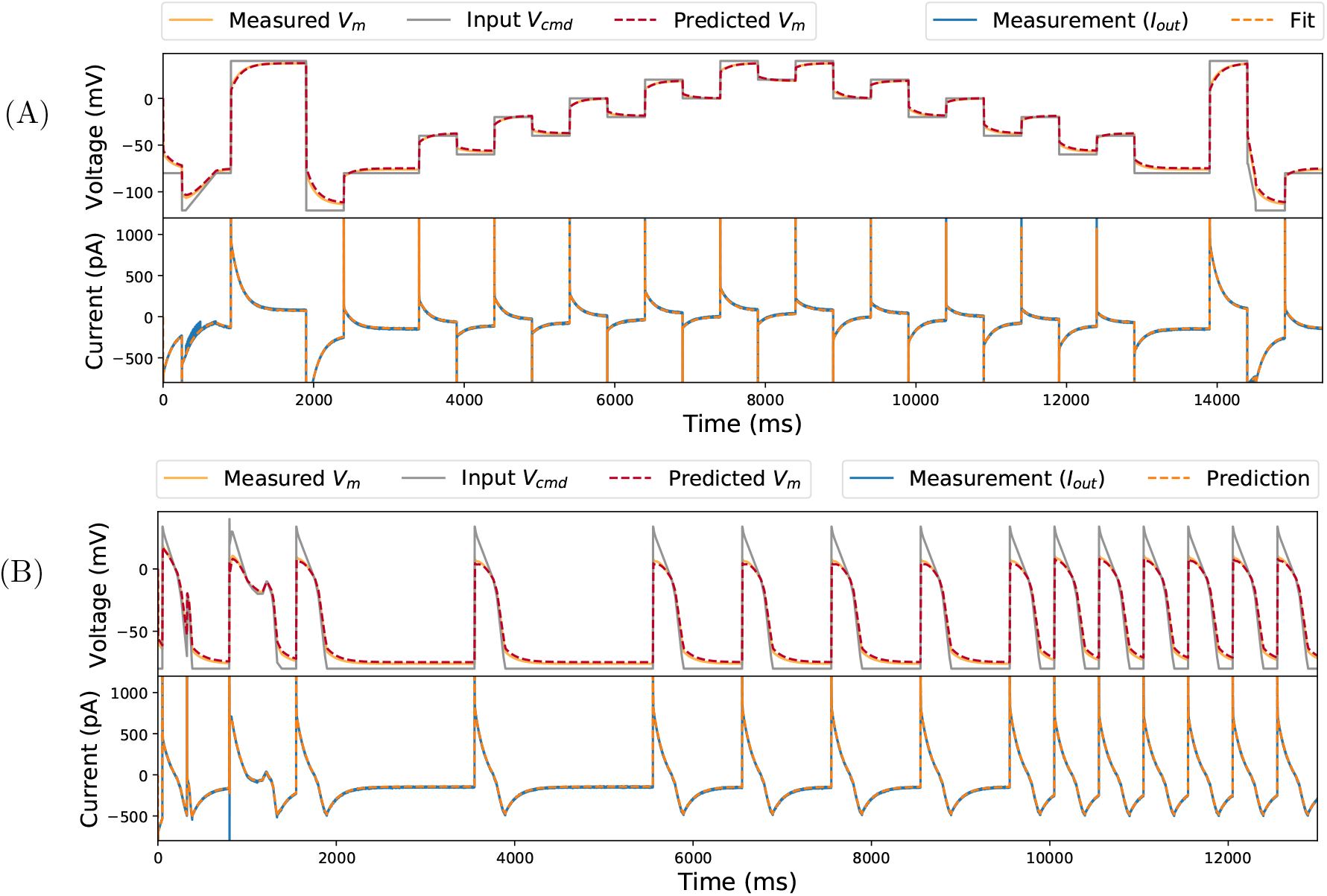
Inferred model simulations and predictions (dashed lines) compared to experimental data (solid lines) from a Type II Model Cell. **(A)** Model calibration with a staircase protocol (grey lines in the top panel), where the model was fitted to only the current recording (blue, solid line in the lower panel). The fitted model was able to predict the membrane voltage, *V*_m_ (orange, solid line) measured using current-clamp. **(B)** Further model validation using an independent voltage-clamp protocol, a series of action potentials (grey lines in the top panel). Again, predictions from the model fitted to the staircase protocol above (dashed lines) are excellent for both the current and the membrane voltage.

Figure 5A shows the fitted model *I*_out_ (bottom, orange dashed line) and its corresponding prediction of the membrane voltage, *V*_m_ (top, red dashed line), compared against the experimental recordings (solid lines). Figure 5B further shows that the fitted model is able to predict measurements under an independent, unseen voltage-clamp protocol — a series of action potential-like waveforms (grey lines in the first panel). Note the excellent predictions for *V*_m_ and its deviation from *V*_cmd_. The series of action potential-like waveforms were concatenated from those in a previous study Lei et al. (2019b), they are composed of linear ramps and steps to permit use on high-throughput machines that cannot clamp to arbitrary waveforms. Magnifications of the protocols are shown later in Figure 7C–F; they include a delayed afterdepolarization (DAD)-like protocol, an early afterdepolarization (EAD)-like protocol, and action-potential-like protocols with different beating frequencies.

Table 2 also shows a comparison of the values of the hardware components (typical manufacturing tolerances for these are ±1 to 2%) in Figure 3, the amplifier’s estimation using a standard test pulse, and the fitted values using the mathematical model. Our model-inferred values are much closer to the component labels than the amplifier estimates because the Type II Model Cell exhibits nonlinear dynamics (as do real cells); whereas the amplifier uses a simple square-wave test pulse and assumes a simple resistor-capacitor model cell (i.e. a Type-I Model Cell) when estimating the parameters. That is, there is a difference between the electrical model cell we attached and the circuit the amplifier is designed to compensate, thus leading to inaccurate estimation of some components. For example, even though we did not apply any voltage offset, *V*_off_, in the experiment, the amplifier incorrectly estimated an offset of −1.2 mV. Proceeding with this amplifier-estimated value would lead to a voltage offset artefact of 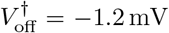 in all recordings. The idea is that using the full model could capture these effects and take them into account when calibrating current kinetic parameters.

**Table 2:** Type II Model Cell parameters (for the components shown in Figure 3). (Row 1) hardware component labels (*V*_off_ is zero as there is no battery/voltage-offset component); (Row 2) Patchmaster amplifier software estimates using a simple test pulse; (Row 3) inferred values from the mathematical model. The mathematical model captures the fact that there are kinetics in the Type II cell and improves on the amplifier’s estimates of the components.

| | $R_k$<br>(MΩ) | $C_k$<br>(pF) | $R_m$<br>(MΩ) | $C_p$<br>(pF) | $C_m$<br>(pF) | $R_s$<br>(MΩ) | $V_{off}$<br>(mV) |
| --- | --- | --- | --- | --- | --- | --- | --- |
| Component label | 100 | 1000 | 500 | 4.7 | 22 | 30 | 0 |
| Patchmaster estimate | — | — | 91.30 | 8.80 | 41.19 | 33.60 | -1.20 |
| Fitted parameters | 94.20 | 1062.69 | 520.70 | 4.85 | 36.38 | 34.87 | 0.20 |

Thus far, we considered a realistic voltage-clamp experiment and developed a detailed mathematical model for such a setting, where imperfect compensations made by the amplifier and imperfect leak current subtraction are included. We then validated this mathematical model via electrical model cell experiments, demonstrating that our model captures the main effects of the voltage-clamp artefacts and amplifier compensations. In the next part of the study, we apply our mathematical model to experimental CHO-hERG1a data recorded previously.

## 4 Application to variability in CHO-hERG1a patch-clamp data

After experimentally validating the mathematical model of the full voltage-clamp experiment with two electrical model cells, it is now applied to experimental data from real cells. Here, a high-throughput dataset from a previous publication is used Lei et al. (2019b). The dataset contains 124 voltage-clamp recordings of the potassium current that flows through hERG (Kv11.1) channels (*I*_Kr_) measured with a staircase protocol and eight other independent protocols. The measurements were performed on CHO cells stably transfected with hERG1a at 25 °C, using the Nanion SyncroPatch 384PE, a 384-well automated patch-clamp platform with temperature control. All the data were leak-corrected (see Eq. (11)), E-4031 subtracted, and passed a semi-automated quality control. The recordings, and parameter values derived from them, showed good agreement with earlier work using manual patch, for details on the methods please see Lei *et al.* Lei et al. (2019b).

In the previous study Lei et al. (2019b), all of the observed variability in the post-processed data was assumed to be due to biological variability in *I*_Kr_ conductance and kinetics, which we term “Hypothesis 1”, and so 124 cell-specific variants/parameterisations of the *I*_Kr_ model were created. Kinetic parameters fitted to cell-specific data using the staircase calibration protocol enabled very good predictions for eight independent validation protocols. However, covariance in the inferred parameters across cells led the authors to speculate about a voltage offset 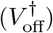 being responsible for much of the variability Lei et al. (2019b).

So an alternative hypothesis is that all cells have the same *I*_Kr_ kinetics (functional properties of the channel proteins), but different maximal conductances (expression levels of the proteins), and that the observed variability in currents is due to differences in the patch-clamp artefacts and compensations for each cell. We term this set of assumptions “Hypothesis 2”. Figure 6 shows a schematic overview of the two hypotheses.

**Figure 6:**
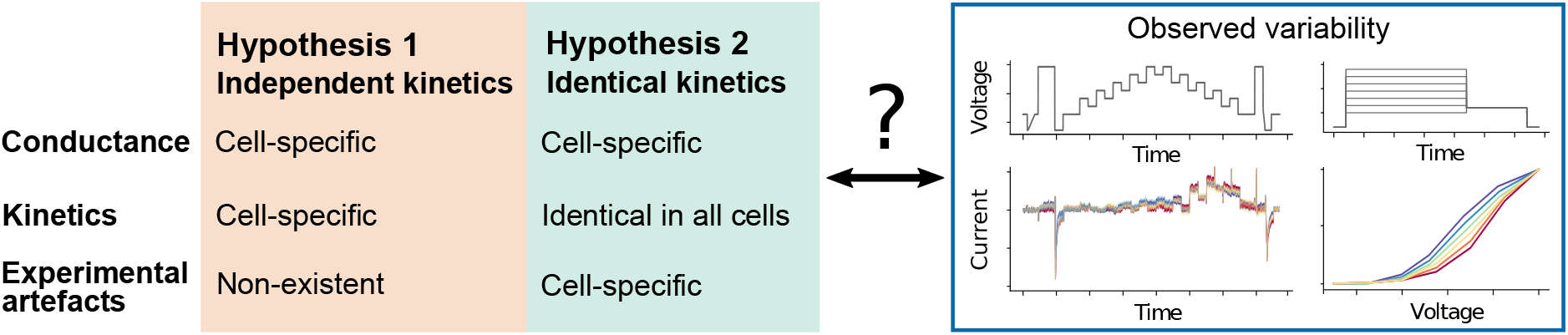
A schematic overview of the two hypotheses we explore. Hypothesis 1 assumes the collected data represent a perfect, idealised voltage-clamp experiment where any experimental artefacts have been perfectly compensated by the amplifier and leak subtraction, so all the observed variability is rooted in biological variability (varying conductance and model kinetics in every cell), we term these ‘independent kinetics models’. Hypothesis 2 assumes that the observed variability is due to differences in the voltage-clamp experimental artefacts, and that all of the cells share identical ion channel kinetics (although the maximum conductance is allowed to vary across cells), we term these ‘identical kinetics models’.

We will now introduce the *I*_Kr_ model and discuss the extent to which patch-clamp artefacts and imperfect compensations can explain the variability in the biological recordings.

### 4.1 A mathematical model of *I*_Kr_

*I*_Kr_ is represented with a model used in previous studies Beattie et al. (2018); Lei et al. (2019b,a); Clerx et al. (2019a). The current is described with two Hodgkin & Huxley-style gating variables (*a* for ‘activation’ and *r* for ‘recovery’ from inactivation) and a standard Ohmic expression,

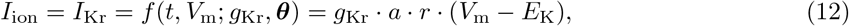

where *g*_Kr_ is the maximal conductance, and *E*_K_ is the reversal potential (Nernst potential) for potassium ions which can be calculated directly from concentrations either side of the membrane using the Nernst equation (Eq. 2 in Lei *et al.* Lei et al. (2019b)). The gates *a* and *r* are governed by the ordinary differential equations

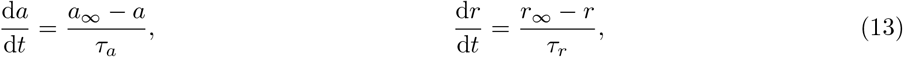

with

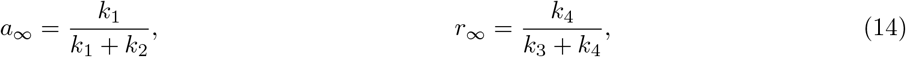

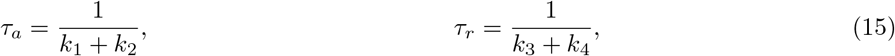

and

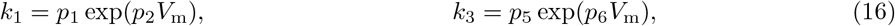

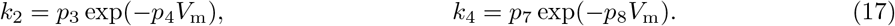

The model has 9 positive parameters: maximal conductance *g*_Kr_ and kinetic parameters ***θ*** = {*p*_1_, *p*_2_, · · ·, *p*_8_} which are optimised to fit the experimental data. Units of the parameters are {pS, s^−1^, V^−1^, s^−1^, · · ·}.

### 4.2 Inference with the full voltage-clamp experiment model

Earlier we saw how the full voltage-clamp experiment model (Eqs. (3–9)) had parameters that could be successfully inferred from data for the electrical model cells. We tested how this inference scheme performs when combined with the *I*_Kr_ model of Eq. (12) in a synthetic data study. All of the parameters could be identified (model parameters *g*_Kr_ and ***θ*** together with artefact parameters *C*_m_, *C*_p_, *R*_s_, 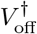 and *g*_leak_) from a single cell recording (full results shown in Supplementary Section S6). We then tested this approach with real experimental data. However, values of the series resistance, *R*_s_, were consistently estimated at the lower bound we had imposed, which we consider unrealistic. This behaviour could be due to imperfections in the representation of *I*_Kr_ by the model (model discrepancy Lei et al. (2020)), as all parameters in the full voltage-clamp experiment model were successfully inferred in the cases of electrical model cells or synthetic *I*_Kr_ data. We therefore propose a simplification of the voltage-clamp experiment model whilst capturing the principal causes of variability.

### 4.3 A simplified voltage-clamp experiment model

Modelling the entire voltage-clamp machine may not be necessary, because the timescales of the components in the system span multiple orders of magnitude; ranging from the order of 0.1 μs (e.g. *τ*_clamp_) to tens of ms (e.g. ‘C-slow’) or many seconds for ion channel gating (e.g. activation of *I*_Kr_ at room temperature Vandenberg et al. (2006)).

For these *I*_Kr_ investigations the two fastest processes, *τ*_clamp_ and *τ*_z_, can be approximated as instantaneous responses. That is, Eqs. (6) & (9) can be approximated as *V*_p_ ≈ *V*_clamp_ and *I*_out_ ≈ *I*_in_, respectively. After analysing the local sensitivity of the voltage-clamp experiment model (Supplementary Section S3), we found that the effects of 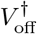 and imperfect *R*_s_ compensation are most apparent in the observed current (on timescales relevant to *I*_Kr_). As a result, we further assume that 1. *τ*_sum_, part of the fast amplifier processes, is instantaneous; 2. the effects of *C*_p_ and *C*_m_ are negligible; and 3. 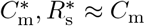, *R*_s_. Finally, the data were leak subtracted, where the leak current parameters (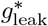 and 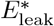) were estimated by fitting Eq. (4) to current during the −120 to −80 mV leak-ramp at the beginning of the measurements, to yield zero current at holding potential Lei et al. (2019b). We allow this leak subtraction to be imperfect by retaining a small residual leak current with parameters 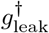 and 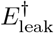.

With these assumptions, Eqs. (3)–(9) become

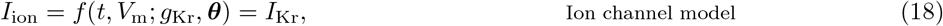

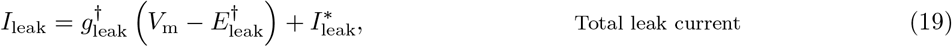

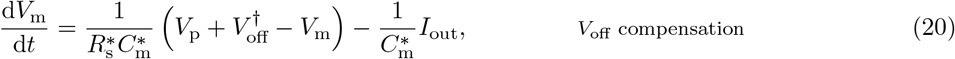

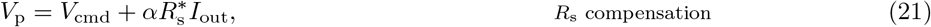

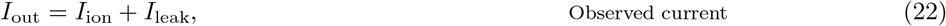

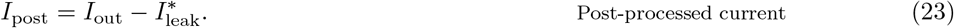

For all symbols refer to Table 1, and 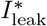 is given by Eq. (10). The effective reversal potential of the residual leak current, 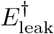, is chosen to be the holding potential (−80 mV) because the primary leak-subtraction (fit of 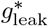 and 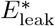) ensured approximately zero current at holding potential. Therefore, we have only two voltage-clamp model parameters (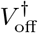 and 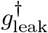) to infer along with the ion current parameters (*g*_Kr_ and ***θ***). Note that the other parameters (*α*, 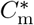, and 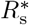) can be obtained from the amplifier settings without performing any inference.

This simplified voltage-clamp experiment model was successfully applied to the model cell experiments (results in Supplementary Section S7) and, like the full model, was able to correct imperfect *R_m_* estimates made by the amplifier.

### 4.4 Optimisation of model parameters

We now present the parameter optimisation schemes used to test each hypothesis.

#### 4.4.1 Hypothesis 1: Cell-specific kinetics with no artefacts

Under Hypothesis 1, any experimental artefacts are perfectly compensated by the amplifier and leak subtraction in post-processing. Any variability in the kinetic parameters obtained in current models fitted to the resulting data would reflect underlying variability in the biological function of the channels. To test this hypothesis, the parameter inference scheme detailed in Lei *et al.* Lei et al. (2019b) was employed. In summary, a Bayesian inference scheme was used which resulted in very narrow posterior distributions. This scheme used some parameter transforms so that the optimisation algorithm Hansen (2006) searches in log-transformed space for certain parameters Clerx et al. (2019a). Here we look for the most likely parameter set under that scheme, which is identical to that given by a least square-error fit. So the log-likelihood of a given parameter set for cell *i* is proportional to

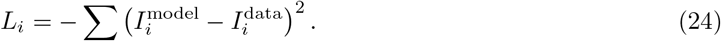

Under this hypothesis, 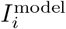 is a function of just conductance *g_i_* and kinetic parameters ***θ***_*i*_, and is given by Eq. (12) while assuming *V*_m_ = *V*_cmd_. So we performed an optimisation to maximise *L_i_* by finding *g_i_* and ***θ***_*i*_ for each cell *i* independently. The resulting models under this hypothesis are termed ‘*independent kinetics models*’.

#### 4.4.2 Hypothesis 2: Identical kinetics for all cells, with cell-specific artefacts

Hypothesis 2 (Figure 6, right) assumes that the observed variability is due to the imperfect voltage-clamp experiments. Under this assumption, models fitted to the data should have the same kinetic parameters, ***θ***\*, across all cells, that is, ***θ***_*i*_ = ***θ***\* for any cell *i*. But there will be a cell-specific *I*_Kr_ conductance, *g_i_*, and different patch-clamp experiment parameters for each cell too, 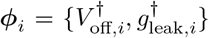. These models are termed ‘*identical kinetics models*’.

To impose the assumption that all *N* cells have the same kinetics, and that the observed variability arises only from the experimental artefacts, the log-likelihood becomes

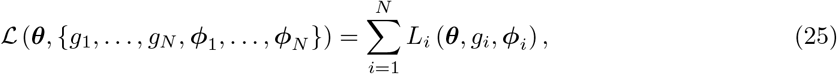

where *L_i_*, is still defined by Eq. (24), the log-likelihood for the *i*^th^ cell. But under this hypothesis, 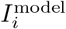 is the post-processed current *I*_post_ in Eq. (23) and hence *L_i_* is a function of the artefact parameters ***ϕ***_*i*_ too.

Optimising 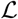 is a high-dimensional optimisation problem, which is computationally expensive. This burden is reduced with a Gibbs-sampling style scheme; breaking the optimisation problem into two: i) optimising the common kinetics parameters, ***θ***; and ii) optimising the cell-specific parameters, {*g_i_*, ***ϕ***_*i*_}_*i*=1,…,*N*_. To evaluate the maximum likelihood of ***θ***, we nest optimisation schemes. That is, for any single estimate of ***θ***, we optimise {*g*_*i*_, ***ϕ***_*i*_} for each cell *i* independently to compute an approximate likelihood for ***θ***. The estimate of ***θ*** is then updated by running a single iteration of the outer optimisation loop, followed again by optimisation to convergence for {*g*_*i*_, ***ϕ***_*i*_} in each cell. This overall scheme converges to the full set of optimal parameters: ***θ***\*, 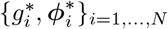. The algorithm is detailed in Supplementary Section S8.

### 4.5 Variability in ion channel kinetics or variability in patch-clamp artefacts?

The performance of the models arising from the two hypotheses is compared next. For Hypothesis 1 the results from Lei *et al.* Lei et al. (2019b) are used, where all the variability was assumed to be the result of kinetic variability (giving 9 parameters× 124 cells = 1116 parameters in total). For Hypothesis 2 we use the results from optimising 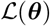 in Eq. (25). A table of optimised parameter values, steady state-voltage relations, and time constants-voltage relations for the new identical kinetics models can be found in Supplementary Section S10.

As in Lei *et al.* Lei et al. (2019b), fits and predictions are quantified using relative root mean square error (RRMSE), defined as the root mean square error between the model simulation and the experimental data, divided by the root mean square distance of the data to a zero current trace:

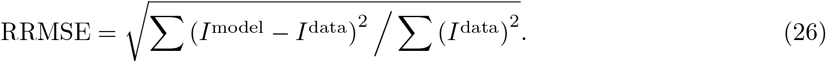

Using this RRMSE quantification, the difference in the absolute size of the current across cells (due to varying conductance) is eliminated such that the scores are comparable between cells.

Figure 7 shows the RRMSE histograms for all 124 cells, for six different protocols, and for the two sets of models. Markers indicate the best (*), median (‡) and 90^th^ percentile (#) RRMSE values for the independent kinetics models (red), and the corresponding raw current traces are shown in the three panels above. The model predictions for the same 3 cells with the identical kinetics model are shown in green. These protocols were used in the previous study Lei et al. (2019b) where they were intended to explore physiologically-relevant predictions for *I*_Kr_ behaviour. The same analysis applied to the remaining 3 protocols is shown in Supplementary Figure S7. The median, 10^th^ percentile, and 90^th^ percentile RRMSE values for each protocol are shown in Supplementary Table S3.

**Figure 7:**
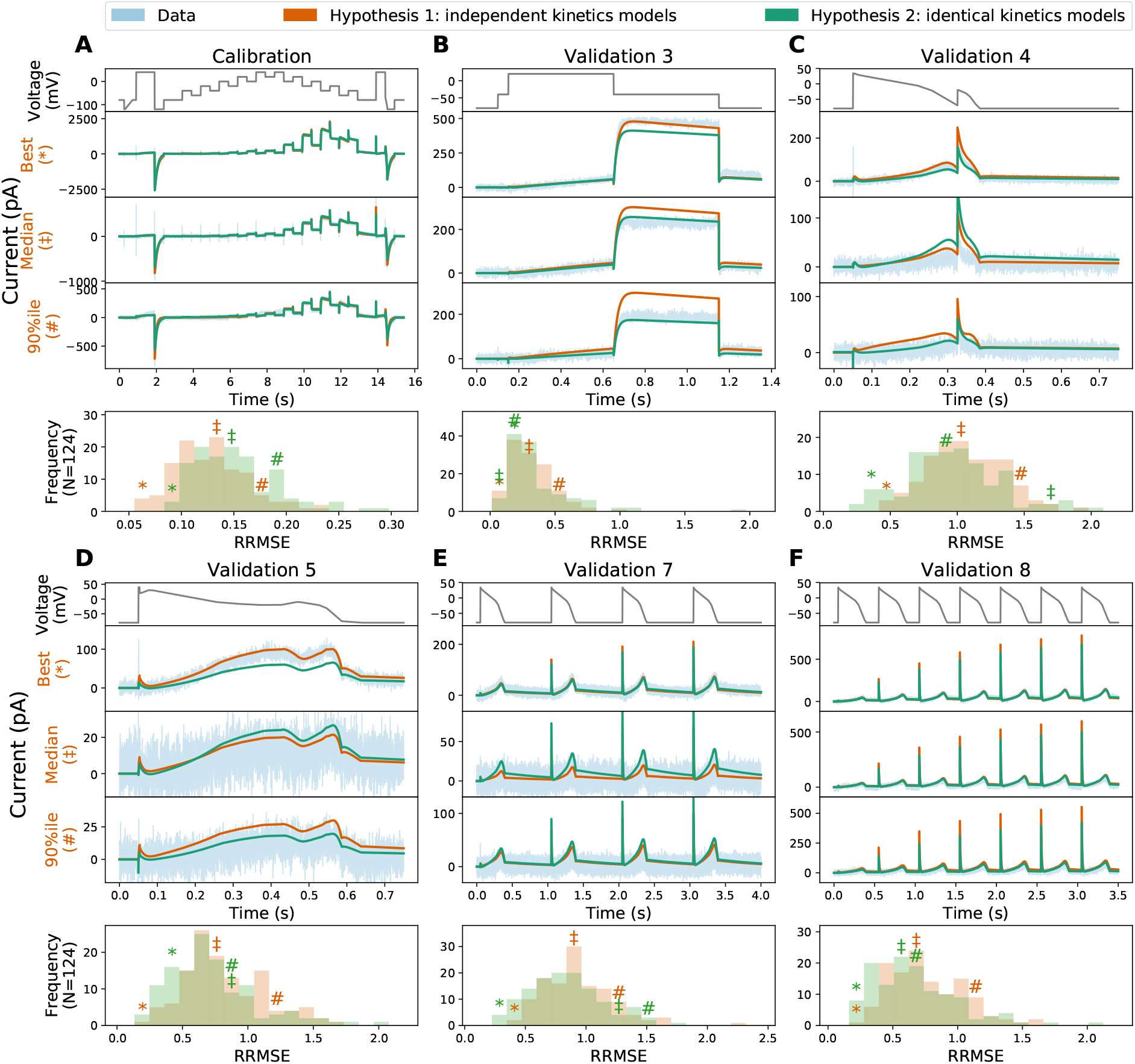
The relative root mean square error (RRMSE) histograms for 6 protocols, comparing the independent kinetics models from Lei *et al.* Lei et al. (2019b) and the identical kinetics models with voltage-clamp artefact. Each histogram represents the same 124 cells with a different protocol and RRMSE each time. Red markers indicate the best (*), median (‡) and 90^th^ percentile (#) RRMSE values for the independent kinetics model; green markers are the same cell prediction from the identical kinetics models. The median, 10^th^ percentile, and 90^th^ percentile of the RRMSE values are shown in Supplementary Table S3. For each protocol, the raw traces for the identical kinetics model (green), the independent kinetics model (red), and data (blue) are shown, with the voltage-clamp above. Note that the currents are shown on different scales for each cell, to reveal the details of the traces. The same analysis applied to the remaining 3 protocols is shown in Supplementary Material. Quantitatively, the two models show a similar RRMSE distribution for each protocol, with a slightly larger error on average in the fit for the identical kinetics (panel A), but a slightly lower error in predictions (panels B-F). In particular, note how the errors seen in two of the cell-specific kinetics predictions (red traces) in panel B are corrected by the identical kinetics model (green).

For the calibration protocol shown in Figure 7A, the RRMSE histogram of independent kinetics models (Hypothesis 1, shown in red) clearly shows lower errors (and hence better fits) than the identical kinetics models (Hypothesis 2, in green). This is expected, as under the independent kinetics hypothesis 9 parameters can be varied per cell (leading to 9 × 124 = 1116 degrees of freedom overall), while under the identical kinetics hypothesis there are only 3 (for 8 + 3 × 124 = 380 degrees of freedom). This increased freedom allows Hypothesis 1 to achieve better fits, but could also lead to *overfitting* Whittaker et al. (2020).

In overfitting, an increased number of degrees-of-freedom leads to improved fits to calibration data, but at the expense of a loss of predictive power on new data sets. Examples in Figure 7B hint that this may be happening. Here, the ‘median quality’ Hypothesis 1 prediction (with 9 cell-specific parameters) over-predicts the peak current under the Validation #3 protocol. In contrast, the Hypothesis 2 prediction (with 3 cell-specific parameters) is much closer to the data, suggesting that the cell-specific Hypothesis 1 parameter set may have varied kinetic parameters to fit details that are better explained by patch artefacts. To summarise such fits across all 124 recordings we can examine the histograms of RRMSE beneath each protocol in Figure 7. Further evidence that overfitting might be occurring under Hypothesis 1 is shown by Figure 7C-F, where the red RRMSE histogram is shifted right, indicating that validation predictions show higher error for Hypothesis 1 than Hypothesis 2. In summary, Hypothesis 2 provides predictions in new situations that overall are at least as good as, if not better than, Hypothesis 1. Coupled with the greatly reduced number of degrees-of-freedom in Hypothesis 2 (736 fewer parameters), this is a strong indication that identical kinetics is the preferred hypothesis for these data.

In the identical kinetics models, all the variability must be explained by the voltage-clamp artefact parameters ***ϕ***_*i*_ and the cell conductance *g*_Kr,*i*_ in Eq. (25). Figure 8 shows the histograms and the pairwise scatter plots of the obtained 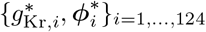. The artefact parameter values are within reasonable ranges: ~ ±5 mV for 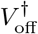 and 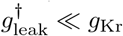 and are not strongly correlated, which lends further credibility to the identical kinetics models (Hypothesis 2).

**Figure 8:**
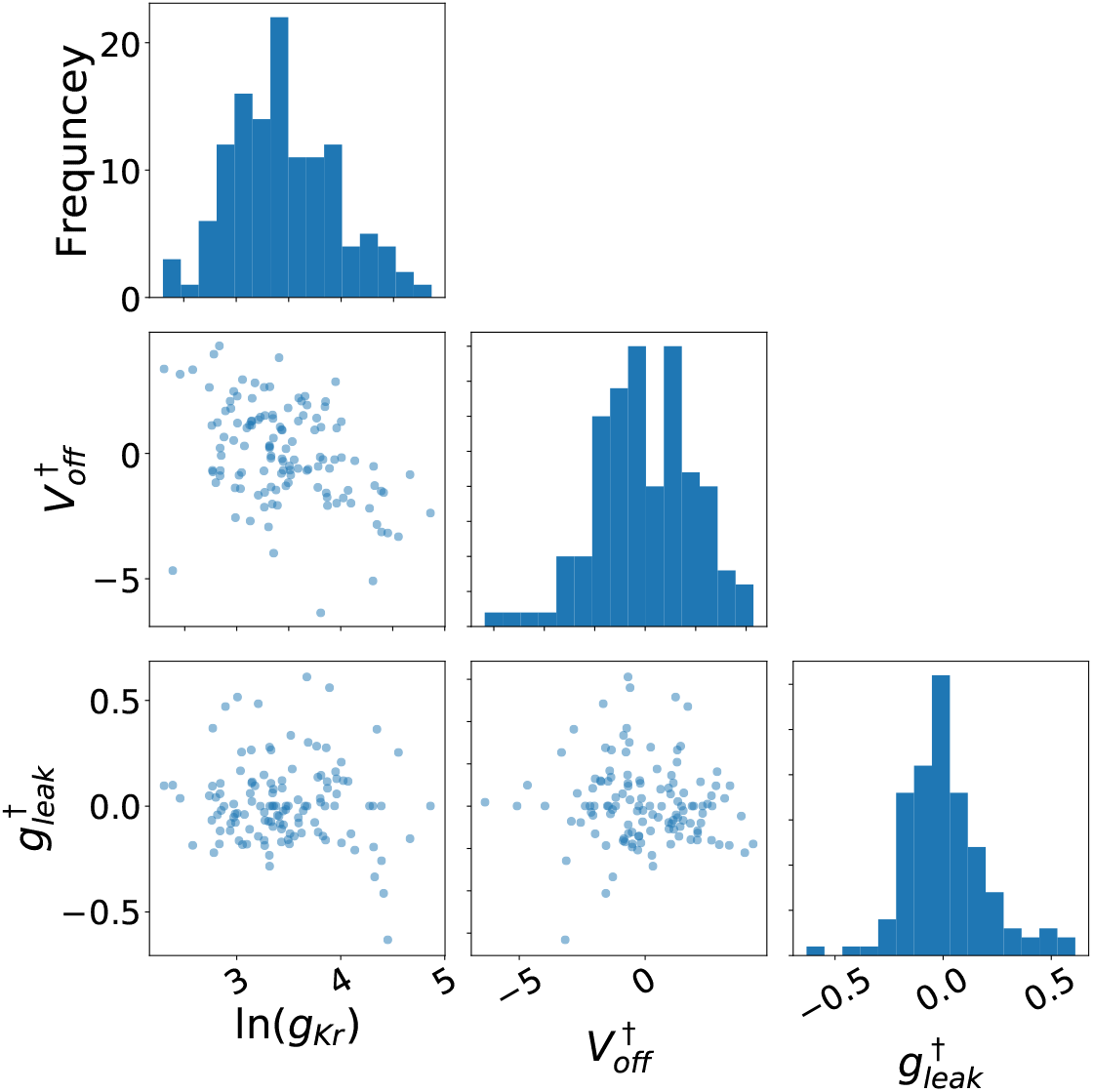
The inferred voltage-clamp artefact parameters across experimental wells. Each of the parameters exhibits a Gaussian-like distribution under this choice of transformation. The artefact parameter values are within reasonable ranges: 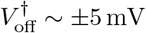 and 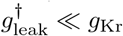.

Finally, one might expect that the mean of the kinetic parameters identified under Hypothesis 1 would be similar to the single set of kinetic parameters identified under Hypothesis 2. Kinetic parameters and model predictions from the two hypotheses are very similar, but not exactly the same (full results in Supplementary Section S10). The difference is due to the unaccounted experimental artefacts in Hypothesis 1 that are included in the (model) kinetic parameters. If Hypothesis 2 is correct, the kinetic parameters derived from it are the more physiologically-relevant.

We also examined a ‘Hypothesis 3’ where conductance, kinetics and artefacts were all cell-specific. Results are omitted for space reasons, but can be summarised as: fits improved slightly (as expected with more parameters) but predictions were of similar quality to Hypothesis 1 and not as good as Hypothesis 2. Figures and code for this analysis can be found in the online repository.

## 5 Discussion

In this paper we have introduced a new mathematical model for voltage-clamp experiments that allows and accounts for experimental artefacts and imperfect compensations of these artefacts, as well as imperfections in leak current subtraction. The model gives insight into the process of how experimental artefacts are introduced and the performance of the compensation procedure. For example, often the largest currents have the highest signal-to-noise ratio and could be interpreted as the ‘cleanest’ recordings. But the model (Eq. (7)) shows that the effect of *R*_s_ in distorting the applied voltage is proportional to the current size. So, for a given series resistance and a fixed percentage compensation (*α*), the bigger the current then the larger this artefact.

We validated the mathematical model through experiments using two types of electrical model cells, where we showed that our mathematical model is able to rectify imperfect amplifier estimations; then applied it to account for artefacts whilst inferring parameters of an *I*_Kr_ model from hERG1a CHO cell data. This is, to our knowledge, the first time a detailed voltage-clamp experiment model has been used for parameter optimisation in ion channel modelling.

Whole-cell patch-clamp data show a high degree of variability Zhou et al. (1998); Vandenberg et al. (2006); Beattie et al. (2018); Lei et al. (2019b,a), with differences in observed current kinetics between experiments. As recordings measure whole-cell ‘macroscopic’ current one might expect variation due to stochastic ‘microscopic’ ion channel opening. But here we have observed currents to be reproducible within a given cell Lei et al. (2019b,a) — that is, the apparent kinetic variability upon protocol repeats in the same cell at different times is much smaller than cell-to-cell variability. We expect the maximum *conductance* of the current to differ between cells, due to varying cell sizes and gene expression levels. Since each CHO cell over-expresses the same channel gene, there is no obvious reason for the current *kinetics* to vary cell-to-cell, especially if the cells are from the same culture and recorded simultaneously under very similar conditions, as they were in our high-throughput data.

Each voltage-clamp experiment has a different cell membrane capacitance, pipette (or well-plate in automated clamp) capacitance, voltage offset, series resistance, and leak current. This led us to propose two competing hypotheses (as shown in Figure 6): Hypothesis 1 is that there are cell-specific kinetics with no artefacts; and Hypothesis 2 is that there are identical kinetics for all cells with cell-specific artefacts. A parameter optimisation technique was developed for Hypothesis 2, so that we were able to optimise all the model parameters at once (cell-specific conductances and measurement artefact parameters, with a single set of kinetic parameters shared by all the cells).

Models based on cell-specific conductances and kinetics (or cell-specific conductances, kinetics and artefacts) included many parameters/degrees-of-freedom. The simpler model of just cell-specific conductances and artefacts made better predictions, on average, over all 124 cells that were analysed. Considering Occam’s razor, this makes the ‘identical kinetics’ hypothesis, in which fewer assumptions are made, the favourable explanation. So it is more plausible that the observed variability in the current traces arose from patch-clamp experimental artefacts, that is imperfect amplifier compensations and imperfect leak current subtraction, than from varying current kinetics. One might expect this result, since the current is carried by thousands of identical ion channel proteins, especially in our over-expression cell line.

It should be noted that these results and this interpretation are specific to our preparation, and cell-cell biological variability in kinetics in general cannot be ruled out. For instance, in native myocytes one might speculate that differing subunit expression and other signalling-related changes in proteins’ states in the membrane (e.g. by PI3K signalling Ballou et al. (2015)) could also confer real biological variability in kinetics from cell to cell. But this study suggests the major factor causing apparent variability in expression system current kinetics will be artefacts introduced by the patch-clamp experimental procedure, and offers an approach to address it.

Determining the origin of variability is particularly important for forward propagation of uncertainty in cardiac modelling. If each ion current biologically exhibited a high-degree of variability in kinetics from cell to cell, then a description of this variability for all currents would be necessary to make predictions at the whole-cell level that accounted for it. However, if the majority of the variability is due to experimental artefacts, then a single set of kinetic parameters could accurately describe the physiology of each current type. If our findings also apply to myocytes, the standard approach to building action potential models with a single set of kinetic parameters for each current is appropriate Ten Tusscher et al. (2004); O’Hara et al. (2011); Groenendaal et al. (2015); Lei et al. (2017), and the observed cell-cell variability in kinetics data is not required to be propagated forward in action potential simulations (as some studies have examined Pathmanathan et al. (2015)). Nevertheless, we may still need an approach like the one demonstrated here to determine unbiased ion channel kinetic parameters to build the most physiologically-relevant action potential models; as opposed to taking the mean of biased recordings which could accidentally include some experimental artefacts within the kinetics models.

Identifying ion current kinetics and separating out experimental artefacts is crucial for many cardiac electrophysiology studies. For example, ion channel mutation studies often conclude with a statement such as “there is a 5–10 mV shift in the half-activation potential” Clerx et al. (2018); Ng et al. (2019). However, given the variability that we observe in patch-clamp data, often the cell to cell variability in half-activation potential can be in the range of 10–15 mV. Therefore, it is important to separate out experimental artefacts from real biological effects. The same principle applies to other cardiac electrophysiology studies, such as drug studies and the basic characterisation of ion channel kinetics.

The findings raise the question of whether a mathematical model of the voltage-clamp experiment could remove the need for amplifier compensations altogether. This approach works well when we have an almost perfect model of the current, as in the electrical model cell cases (see Figure 5). But we do not recommend it in general, because unlike the reduced model, the full voltage-clamp experiment model (Eqs. (3)–(9)) with an *I*_Kr_ model appeared to produce spurious fits to real cell data. This behaviour is possibly due to *I*_Kr_ model discrepancy — extra artefact components in the full model may attempt to ‘mop up’ real *I*_Kr_ current which the approximations in this simple *I*_Kr_ model do not explain. We envision that a more detailed *I*_Kr_ model, or techniques for accounting for discrepancy Lei et al. (2020), could allow us to infer parameters for the full patch-clamp model. Yet our simplified voltage-clamp model (Eqs. (18)–(23)) appears to account for imperfect amplifier compensation even with an inevitably imperfect mathematical model of *I*_Kr_ kinetics.

In this study we have demonstrated that a reduced voltage-clamp experiment model can unify the kinetics of *I*_Kr_ measured across many cells. Our model represents the voltage-clamp experimental set-up and could be applied to any voltage-clamp data gathered with a standard patch-clamp amplifier. Our full model of the voltage-clamp experiment could be particularly useful for currents like the fast sodium current (*I*_Na_) where the time constants are similar to those in the capacitance artefacts.

Finally, it should be possible to generalise our method to current-clamp experiments by introducing an extra feedback circuit to the full voltage-clamp model for standard patch-clamp amplifiers HEKA Elektronik Dr. Schulze GmbH (2007–2018); Axon Instruments Inc. (1997–1999). In current-clamp mode, the leak current correction is usually performed during the measurement, therefore Eq. (23) needs to be incorporated into Eq. (8). However, note that conventional microelectrode amplifiers, such as the Axoclamp and the Axoprobe, have a different headstage design and a re-analysis of their circuits to build a similar model would need to be undertaken Axon Instruments Inc. (1997–1999).

## 6 Conclusions

A mathematical model was derived to describe the entire voltage-clamp experiment including artefacts, imperfect amplifier compensations, and imperfect leak current subtraction. Using this model, variability in a set of experimental observations could be explained either by varying current properties or varying measurement artefacts. After comparing model predictions in each case, the results suggest that most of the observed variability in a set of expression system patch-clamp data measured in high-throughput under the same conditions is caused by experimental artefacts. These varying experimental artefacts can be compensated in post-processing by fitting the mathematical model for the patch-clamp experiment at the same time as fitting ion channel kinetics. This study raises questions for the biological significance of any cell-cell variability in macroscopic ion channel kinetics, and provides for better correction of artefacts in patch-clamp data.

## Data access

All codes and data are freely available at https://github.com/CardiacModelling/VoltageClampModel.

## Author contributions

CLL, DGW & TPdB built the model cells and carried out the experiments. CLL, MC & GRM derived the mathematical models. CLL wrote code for and performed the simulations and data analysis, and generated the results figures. CLL, MC, DJG, TPdB and GRM conceived of and designed the study. All authors drafted and approved the final version of the manuscript.

## Acknowledgements

We thank Alan Fabbri of UMC Utrecht for assistance with the model cell measurements.

## References

Altomare, C., Bartolucci, C., Sala, L., Bernardi, J., Mostacciuolo, G., Rocchetti, M., Severi, S., Zaza, A., 2015. IKr impact on repolarization and its variability assessed by dynamic clamp. Circulation: Arrhythmia and Electrophysiology 8, 1265–1275.

Annecchino, L.A., Schultz, S.R., 2018. Progress in automating patch clamp cellular physiology. Brain and Neuroscience Advances 2.

Axon Instruments Inc., 1997–1999. Axopatch 200B patch clamp theory and operation. https://www.autom8.com/wp-content/uploads/2016/07/Axopatch-200B.pdf. Accessed: 2020-02-29.

Ballou, L.M., Lin, R.Z., Cohen, I.S., 2015. Control of cardiac repolarization by phosphoinositide 3-kinase signaling to ion channels. Circulation research 116, 127–137.

Beattie, K.A., Hill, A.P., Bardenet, R., Cui, Y., Vandenberg, J.I., Gavaghan, D.J., De Boer, T.P., Mirams, G.R., 2018. Sinusoidal voltage protocols for rapid characterisation of ion channel kinetics. The Journal of Physiology 596, 1813–1828.

Bekkers, J., Richerson, G., Stevens, C., 1990. Origin of variability in quantal size in cultured hippocampal neurons and hippocampal slices. Proceedings of the National Academy of Sciences 87, 5359–5362.

Clerx, M., Beattie, K.A., Gavaghan, D.J., Mirams, G.R., 2019a. Four ways to fit an ion channel model. Biophyical Journal 117, 2420–2437.

Clerx, M., Collins, P., de Lange, E., Volders, P.G.A., 2016. Myokit: A simple interface to cardiac cellular electrophysiology. Progress in Biophysics and Molecular Biology 120, 100–114.

Clerx, M., Heijman, J., Collins, P., Volders, P.G., 2018. Predicting changes to INa from missense mutations in human SCN5A. Scientific Reports 8, 12797.

Clerx, M., Robinson, M., Lambert, B., Lei, C.L., Ghosh, S., Mirams, G.R., Gavaghan, D.J., 2019b. Probabilistic inference on noisy time series (PINTS). Journal of Open Research Software 7, 23.

Feigenspan, A., Dedek, K., Schlich, K., Weiler, R., Thanos, S., 2010. Expression and biophysical characterization of voltage-gated sodium channels in axons and growth cones of the regenerating optic nerve. Investigative Ophthalmology & Visual Science 51, 1789–1799.

Finkel, A., Wittel, A., Yang, N., Handran, S., Hughes, J., Costantin, J., 2006. Population patch clamp improves data consistency and success rates in the measurement of ionic currents. Journal of Biomolecular Screening 11, 488–496.

Golowasch, J., 2014. Ionic current variability and functional stability in the nervous system. Bioscience 64, 570–580.

Groenendaal, W., Ortega, F.A., Kherlopian, A.R., Zygmunt, A.C., Krogh-Madsen, T., Christini, D.J., 2015. Cell-specific cardiac electrophysiology models. PLOS Computational Biology 11, e1004242.

Hansen, N., 2006. The CMA Evolution Strategy: A comparing review, in: Towards a New Evolutionary Computation: Advances in the Estimation of Distribution Algorithms. Springer Berlin Heidelberg, Berlin, Heidelberg, pp. 75–102.

HEKA Elektronik Dr. Schulze GmbH, 2005–2016. PATCHMASTER multi-channel data acquisition software reference manual 2×90.2. https://www.heka.com/downloads/software/manual/m_patchmaster.pdf. Accessed: 2020-02-29.

HEKA Elektronik Dr. Schulze GmbH, 2007–2018. EPC 10 USB hardware manual version 2.8. http://www.heka.com/downloads/hardware/manual/m_epc10.pdf. Accessed: 2020-02-29.

Lei, C.L., Clerx, M., Beattie, K.A., Melgari, D., Hancox, J.C., Gavaghan, D.J., Polonchuk, L., Wang, K., Mirams, G.R., 2019a. Rapid characterisation of hERG channel kinetics II: temperature dependence. Biophysical Journal 117, 2455–2470.

Lei, C.L., Clerx, M., Gavaghan, D.J., Polonchuk, L., Mirams, G.R., Wang, K., 2019b. Rapid characterisation of hERG channel kinetics I: using an automated high-throughput system. Biophysical Journal 117, 2438–2454.

Lei, C.L., Ghosh, S., Whittaker, D.G., et al., 2020. Considering discrepancy when calibrating a mechanistic electrophysiology model. Philosophical Transactions of the Royal Society A this volume.

Lei, C.L., Wang, K., Clerx, M., Johnstone, R.H., Hortigon-Vinagre, M.P., Zamora, V., Allan, A., Smith, G.L., Gavaghan, D.J., Mirams, G.R., Polonchuk, L., 2017. Tailoring mathematical models to stem-cell derived cardiomyocyte lines can improve predictions of drug-induced changes to their electrophysiology. Frontiers in Physiology 8.

Li, Z., Ridder, B.J., Han, X., Wu, W.W., Sheng, J., Tran, P.N., Wu, M., Randolph, A., Johnstone, R.H., Mirams, G.R., et al., 2019. Assessment of an in silico mechanistic model for proarrhythmia risk prediction under the CiPA initiative. Clinical Pharmacology & Therapeutics 105, 466–475.

Marty, A., Neher, E., 1995. Tight-seal whole-cell recording, in: Sakmann, B., Neher, E. (Eds.), Single-Channel Recording. Springer US, Boston, MA. chapter 2, 2 edition. pp. 31–52.

Mirams, G.R., Davies, M.R., Cui, Y., Kohl, P., Noble, D., 2012. Application of cardiac electrophysiology simulations to pro-arrhythmic safety testing. British Journal of Pharmacology 167, 932–945.

Mirams, G.R., Pathmanathan, P., Gray, R.A., Challenor, P., Clayton, R.H., 2016. Uncertainty and variability in computational and mathematical models of cardiac physiology. The Journal of Physiology 594, 6833–6847.

Moore, J.W., Hines, M., Harris, E.M., 1984. Compensation for resistance in series with excitable membranes. Biophysical Journal 46, 507–14.

Neher, E., 1992. [6] Correction for liquid junction potentials in patch clamp experiments, in: Methods in Enzymology. Academic Press. volume 207 of Ion Channels, pp. 123–131.

Neher, E., 1995. Voltage offsets in patch-clamp experiments, in: Sakmann, B., Neher, E. (Eds.), Single-Channel Recording. Springer US, Boston, MA. chapter 6, 2 edition. pp. 147–153.

Ng, C.A., Perry, M.D., Liang, W., Smith, N.J., Foo, B., Shrier, A., Lukacs, G.L., Hill, A.P., Vandenberg, J.I., 2019. High-throughput phenotyping of heteromeric human ether-à-go-go-related gene potassium channel variants can discriminate pathogenic from rare benign variants. Heart Rhythm.

Niederer, S.A., Lumens, J., Trayanova, N.A., 2018. Computational models in cardiology. Nature Reviews Cardiology 16, 100–111.

O’Hara, T., Virág, L., Varró, A., Rudy, Y., 2011. Simulation of the undiseased human cardiac ventricular action potential: model formulation and experimental validation. PLOS Computational Biology 7, e1002061.

Pathmanathan, P., Shotwell, M.S., Gavaghan, D.J., Cordeiro, J.M., Gray, R.A., 2015. Uncertainty quantification of fast sodium current steady-state inactivation for multi-scale models of cardiac electrophysiology. Progress in Biophysics & Molecular Biology 117, 4–18.

Raba, A.E., Cordeiro, J.M., Antzelevitch, C., Beaumont, J., 2013. Extending the conditions of application of an inversion of the hodgkin–huxley gating model. Bulletin of mathematical biology 75, 752–773.

Sakakibara, Y., Furukawa, T., Singer, D.H., Jia, H., Backer, C.L., Arentzen, C.E., Wasserstrom, J.A., 1993. Sodium current in isolated human ventricular myocytes. American Journal of Physiology-Heart and Circulatory Physiology 265, H1301–H1309.

Santillo, S., Moriello, A.S., Di Maio, V., 2014. Electrophysiological variability in the SH-SY5Y cellular line. General Physiology and Biophysics 33, 121–129.

Satoh, H., Delbridge, L., Blatter, L.A., Bers, D.M., 1996. Surface: volume relationship in cardiac myocytes studied with confocal microscopy and membrane capacitance measurements: species-dependence and developmental effects. Biophysical Journal 70, 1494–1504.

Schmitt, B.M., Koepsell, H., 2002. An improved method for real-time monitoring of membrane capacitance in Xenopus laevis oocytes. Biophysical Journal 82, 1345–1357.

Sherman, A.J., Shrier, A., Cooper, E., 1999. Series Resistance Compensation for Whole-Cell Patch-Clamp Studies Using a Membrane State Estimator. Biophysical Journal 77, 2590–2601.

Sigworth, F., 1995a. Design of the EPC-9, a computer-controlled patch-clamp amplifier. 1. Hardware. Journal of Neuroscience Methods 56, 195–202.

Sigworth, F., 1995b. Electronic design of the patch clamp, in: Sakmann, B., Neher, E. (Eds.), Single-Channel Recording. Springer US, Boston, MA. chapter 6, 2 edition. pp. 95–127.

Sigworth, F., Affolter, H., Neher, E., 1995. Design of the EPC-9, a computer-controlled patch-clamp amplifier. 2. Software. Journal of Neuroscience Methods 56, 203–215.

Strickholm, A., 1995. A single electrode voltage, current-and patch-clamp amplifier with complete stable series resistance compensation. Journal of Neuroscience Methods 61, 53–66.

Ten Tusscher, K., Noble, D., Noble, P.J., Panfilov, A.V., 2004. A model for human ventricular tissue. American Journal of Physiology-Heart and Circulatory Physiology 286, H1573–H1589.

US National Research Council, 2012. Assessing the reliability of complex models: mathematical and statistical foundations of verification, validation, and uncertainty quantification. National Academies Press.

Vandenberg, J.I., Varghese, A., Lu, Y., Bursill, J.A., Mahaut-Smith, M.P., Huang, C.L.H., 2006. Temperature dependence of human ether-a-go-go-related gene K+ currents. American Journal of Physiology-Cell Physiology 291, C165–75.

Weerakoon, P., Culurciello, E., Klemic, K.G., Sigworth, F.J., 2009. An Integrated Patch-Clamp Potentiostat With Electrode Compensation. IEEE Transactions on Biomedical Circuits and Systems 3, 117–125.

Weerakoon, P., Culurciello, E., Yang, Y., Santos-Sacchi, J., Kindlmann, P.J., Sigworth, F.J., 2010. Patch-clamp amplifiers on a chip. Journal of Neuroscience Methods 192, 187–92.

Whittaker, D.G., Clerx, M., Lei, C.L., Christini, D.J., Mirams, G.R., 2020. Calibration of ionic and cellular cardiac electrophysiology models. WIREs Systems Biology and Medicine, e1482https://onlinelibrary.wiley.com/doi/pdf/10.1002/wsbm.1482.

Zhou, Z., Gong, Q., Ye, B., Fan, Z., Makielski, J.C., Robertson, G.A., January, C.T., 1998. Properties of HERG channels stably expressed in HEK293 cells studied at physiological temperature. Biophysical Journal 74, 230–241.

