## Supplementary Information for "Accounting for variability in ion current recordings using a mathematical model of artefacts in voltage-clamp experiments"

#### S1 Ideal voltage-clamp model

Figure S1 shows an idealised voltage-clamp experiment, a schematic set-up (left) and its equivalent circuit (right), where the cell is connected directly to an ammeter which records the current of interest, $I_{\text{ion}}$, while clamping at the command voltage, $V_{\text{cmd}}$. That is, it assumes

$$\text{(membrane voltage) } V_{\text{m}} = V_{\text{cmd}} \text{ (command voltage)},$$
$$\text{(measured/observable current) } I_{\text{out}} = I_{\text{ion}} \text{ (current of interest)}.$$

Most patch-clamp and modelling studies use these idealised assumptions when analysing voltage-clamp experiments or recordings.

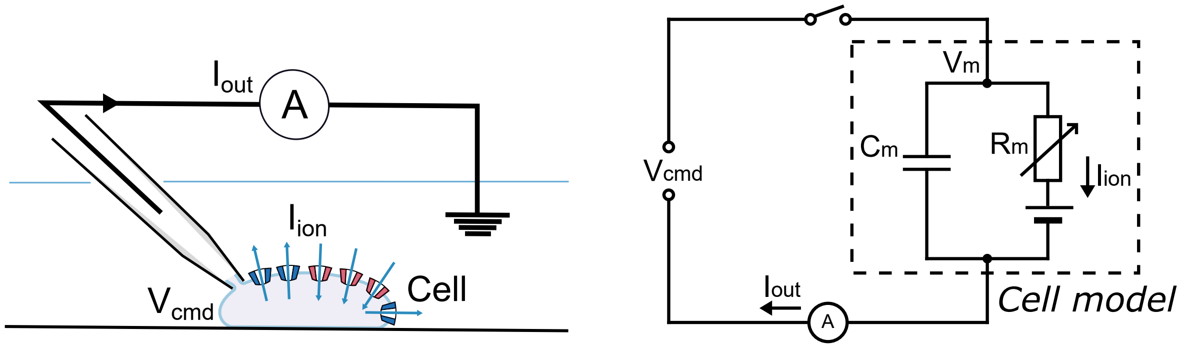


Figure S1: An ideal voltage-clamp experiment schematic set-up **(left)** and its equivalent circuit **(right)**. The membrane voltage, $V_{\text{m}}$, is assumed to be exactly the same as the clamped command voltage, $V_{\text{cmd}}$, and the measured current, $I_{\text{out}}$, is purely the current conducted by the ion channel, $I_{\text{ion}}$. Despite being unrealistic, this is one of the most common assumptions that modelling studies makes when analysing voltage-clamp experiments or recordings (i.e. assuming that patch clamp compensations give rise to the above situation).

### S2 A detailed derivation of the voltage clamp experiment model

A description of all of the symbols and parameters used in this derivation can be found in Table 1 in the main text.

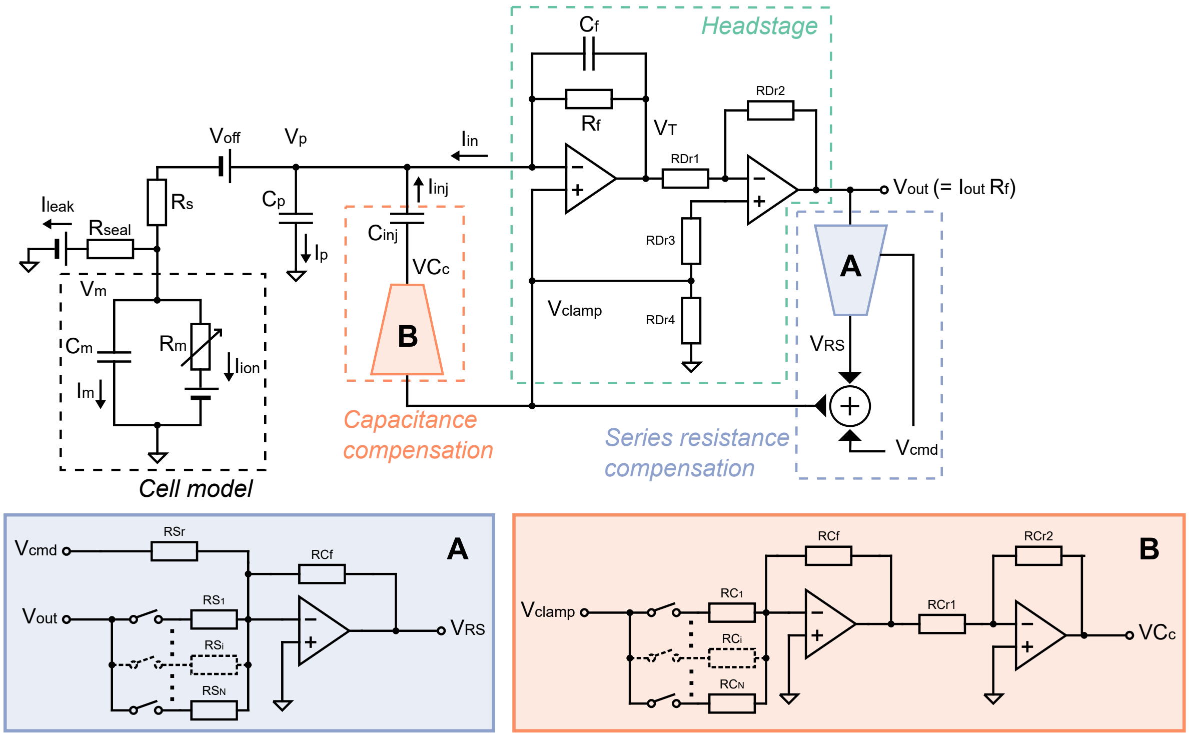


Figure S2: A realistic voltage-clamp experiment equivalent circuit. This includes undesired factors such as voltage offset, series resistance between the electrode and the cell, cell capacitance, pipette capacitance, and leakage current, which can introduce artefacts to the recordings. The circuit also includes the components within a typical amplifier that are designed to compensate the artefacts. The blue **(A)** and orange **(B)** components are two idealised multiplying digital-to-analogue converters (mDACs) or equivalent circuits that control the amount of compensation, which we assume work exactly as intended, so that only their overall effects need to be modelled.

First of all, at the electrode-membrane junction in Figure S2, by applying Kirchhoff's current law, we have

$$I_{\mathrm{m}} = I_{\mathrm{in}} - I_{\mathrm{p}} - I_{\mathrm{leak}} - I_{\mathrm{ion}}, \tag{S2.1}$$

where

$$C_{\mathrm{p}} \frac{\mathrm{d}V_{\mathrm{p}}}{\mathrm{d}t} = I_{\mathrm{p}}, \tag{S2.2}$$

and

$$C_{\mathrm{m}} \frac{\mathrm{d}V_{\mathrm{m}}}{\mathrm{d}t} = I_{\mathrm{m}}. \tag{S2.3}$$

Therefore

$$C_{\mathrm{m}} \frac{\mathrm{d}V_{\mathrm{m}}}{\mathrm{d}t} = I_{\mathrm{in}} - C_{\mathrm{p}} \frac{\mathrm{d}V_{\mathrm{p}}}{\mathrm{d}t} - I_{\mathrm{leak}} - I_{\mathrm{ion}}, \tag{S2.4}$$

$$I_{\mathrm{in}} = I_{\mathrm{ion}} + I_{\mathrm{leak}} + C_{\mathrm{m}} \frac{\mathrm{d}V_{\mathrm{m}}}{\mathrm{d}t} + C_{\mathrm{p}} \frac{\mathrm{d}V_{\mathrm{p}}}{\mathrm{d}t}. \tag{S2.5}$$

This shows that $I_{\mathrm{in}}$ is 'contaminated' by a leak current, a C-Slow ($C_{\mathrm{m}}$) term, and a C-Fast ($C_{\mathrm{p}}$) term. Here, $I_{\mathrm{in}}$ is observed via $V_{\mathrm{out}}$ using the transimpedance amplifier within the Headstage (green box in Fig. S2), a feedback system that converts an input current to a voltage output, which is low-pass filtered by the transconductor time constant $\tau_z = R_f C_f$. We then have

$$V_{\text{out}} + \tau_z \frac{\mathrm{d}V_{\text{out}}}{\mathrm{d}t} = I_{\text{in}} R_f, \tag{S2.6}$$

and since $V_{\text{out}} = I_{\text{out}} R_f$, we have

$$I_{\text{out}} + \tau_z \frac{\mathrm{d}I_{\text{out}}}{\mathrm{d}t} = I_{\text{in}}. \tag{S2.7}$$

Therefore, our final observed current $I_{\text{out}}$ is given by

$$\frac{\mathrm{d}I_{\text{out}}}{\mathrm{d}t} = \frac{I_{\text{in}} - I_{\text{out}}}{\tau_z}. \tag{S2.8}$$

Furthermore, most of the current measured using voltage-clamp depends on $V_m$ which we try to control through the command voltage, $V_{\text{cmd}}$. By analysing the voltage drop across the series resistance, $R_s$, we have

$$(V_p + V_{\text{off}}) - V_m = R_s (I_{\text{in}} - I_p), \tag{S2.9}$$

and together with Eq. (S2.4), we get

$$C_m \frac{\mathrm{d}V_m}{\mathrm{d}t} = \frac{V_p + V_{\text{off}} - V_m}{R_s} - I_{\text{ion}} - I_{\text{leak}}. \tag{S2.10}$$

This is more usually written as

$$\frac{\mathrm{d}V_m}{\mathrm{d}t} = \frac{1}{\tau_a}(V_p + V_{\text{off}} - V_m) - \frac{1}{C_m}(I_{\text{ion}} - I_{\text{leak}}), \tag{S2.11}$$

where $\tau_a = R_s C_m$, and $V_{\text{off}}$ is the voltage offset. From Eq. (S2.11) we can see that as $R_s \to 0$, $V_m \to V_p$ (with an offset $V_{\text{off}}$) instantly; whilst, as $R_s \to \infty$, $V_m$ behaves independently of $V_p$ and is determined by the membrane ion channels as if an isolated cell. The pipette voltage $V_p$ is controlled by the command voltage $V_{\text{cmd}}$, which are delayed by the electrical components,

$$\frac{\mathrm{d}V_p}{\mathrm{d}t} = \frac{1}{\tau_{\text{clamp}}}(V_{\text{clamp}} - V_p), \tag{S2.12}$$

and

$$V_{\text{clamp}} = V_{\text{cmd}}. \tag{S2.13}$$

Finally, we model a linear-in-voltage leak current of the form

$$I_{\text{leak}} = g_{\text{leak}} (V_m - E_{\text{leak}}). \tag{S2.14}$$

If this is a clean setup with no other leaks apart from a non-selective ion current leak through the pipette-cell seal, the parameters of this current would be $g_{\text{leak}} = 1/R_{\text{seal}}$ and $E_{\text{leak}} = 0$. Note that this is usually compensated/subtracted as part of the post-processing, and hence we denote the parameters as $g^{\dagger}_{\text{leak}}$ and $E^{\dagger}_{\text{leak}}$ to reflect the fact that they are the error of the estimate (residual).

After analysing all the undesired artefacts we mathematically model how modern patch amplifiers typically compensate for them Weerakoon et al. (2009); Axon Instruments Inc. (1997–1999); HEKA Elektronik Dr. Schulze GmbH (2007–2018). Firstly, the voltage offset is usually estimated and compensated *prior* to adding the cell to the system, either with an automated correction estimated using *software control* or by applying manually a voltage offset such that it gives zero current when clamped at zero voltage, so the compensation circuit is not shown in our patch clamp equivalent circuits Neher (1995); Sigworth et al. (1995). The major source of voltage offset may be the liquid junction potential, a potential difference of $\sim 2 - 12$ mV which develops when the pipette-filling solution is different from the bath solution Neher (1992). The adjustment is usually done by adding the machine estimated voltage offset $V^*_{\text{off}}$ to $V_{\text{cmd}}$. We can write the error in the estimate of voltage offset $V^{\dagger}_{\text{off}}$ as

$$V^{\dagger}_{\text{off}} = V_{\text{off}} - V^*_{\text{off}}. \tag{S2.15}$$

We then simply need to replace all instances of $V_{\text{off}}$ in the equations above with $V^{\dagger}_{\text{off}}$ to describe the effect of imperfect voltage offset compensation, and $V^{\dagger}_{\text{off}}$ is assumed to be $\mathcal{O}(10)$ mV.

Secondly, to compensate the effect of the parasitic capacitance at the electrode, an additional current $I_{\text{inj}}$ is injected at the electrode to compensate for the current drawn by the parasitic capacitance. By analysing the capacitance compensation part (orange box in Figure S2), we get

$$I_{\text{inj}} = C_{\text{inj}} \frac{\mathrm{d}V_{\text{clamp}}}{\mathrm{d}t}, \tag{S2.16}$$

where $C_{\text{inj}}$ is the amplifier's estimate of the parasitic capacitance $C_{\text{p}}$. Then Eq. (S2.1) becomes

$$I_{\text{m}} = I_{\text{in}} + I_{\text{inj}} - I_{\text{p}} - I_{\text{leak}} - I_{\text{ion}}, \tag{S2.17}$$

and Eq. (S2.5) becomes

$$I_{\text{in}} = I_{\text{ion}} + I_{\text{leak}} + C_{\text{m}}\frac{\text{d}V_{\text{m}}}{\text{d}t} + \left(C_{\text{p}}\frac{\text{d}V_{\text{p}}}{\text{d}t} - C_{\text{inj}}\frac{\text{d}V_{\text{clamp}}}{\text{d}t}\right). \tag{S2.18}$$

This is usually known as 'C-Fast' compensation.

Thirdly, we need to consider compensation for the cell membrane capacitance $C_{\text{m}}$. Usually the effect of $C_{\text{m}}$ is reduced by a hardware 'C-Slow' compensation, using a similar circuit to the 'C-Fast' compensation discussed above Sigworth (1995); Sigworth et al. (1995). However, since the value of $C_{\text{m}}$ can reach 100 pF in some cell types, and capacitor sizes can be limited, the 'C-Slow' compensation is sometimes performed as a post-processing step by the amplifier control software rather than using built-in amplifier hardware Weerakoon et al. (2010). In either case, the full capacitance compensation can be written as

$$I_{\text{in}} = I_{\text{ion}} + I_{\text{leak}} + \left(C_{\text{m}}\frac{\text{d}V_{\text{m}}}{\text{d}t} - C_{\text{m}}^{*}\frac{\text{d}V_{\text{clamp}}}{\text{d}t}\right) + \left(C_{\text{p}}\frac{\text{d}V_{\text{p}}}{\text{d}t} - C_{\text{inj}}\frac{\text{d}V_{\text{clamp}}}{\text{d}t}\right), \tag{S2.19}$$

where $C_{\text{m}}^{*}$ is the amplifier (or user's) estimate of the membrane capacitance $C_{\text{m}}$.

Finally, a series resistance compensation Moore et al. (1984); Strickholm (1995); Sigworth (1995); Sigworth et al. (1995); Sherman et al. (1999); Weerakoon et al. (2009) is implemented to reduce the effect of $\tau_{\text{a}}$ caused by $R_{\text{s}}$ in Eq. (S2.11). By analysing the series resistance compensation part (blue box in Figure S2), we have

$$\frac{\text{d}V_{\text{clamp}}}{\text{d}t} = \frac{1}{\tau_{\text{sum}}}\big((V_{\text{cmd}} + \alpha R_{\text{s}}^{*} I_{\text{out}}) - V_{\text{clamp}}\big), \tag{S2.20}$$

where $R_{\text{s}}^{*}$ is the machine estimation of the series resistance $R_{\text{s}}$, and $\alpha$ is the fraction of $R_{\text{s}}^{*}$ to be compensated (proportion of series resistance compensation).

A summary of the voltage-clamp experiment model equations is shown in Eqs. (3)–(9) in the main text.

#### S3 A sensitivity analysis of the voltage-clamp experiment model

We perform a simple local sensitivity analysis of the voltage-clamp experiment model to study the behaviour of the model. We look at the sensitivity of the 4 parameters, $C_\mathrm{m}^*$, $R_\mathrm{s}^*$, $C_\mathrm{p}^\dagger$ and $V_\mathrm{off}^\dagger$, in the output/observed current. The values that we used are $C_\mathrm{m}^* = 10\,\mathrm{pF}$, $R_\mathrm{s}^* = 5\,\mathrm{M\Omega}$, $C_\mathrm{p}^\dagger = 0\,\mathrm{pF}$ and $V_\mathrm{off}^\dagger = 0\,\mathrm{mV}$. We assume all settings ($C_\mathrm{m}$, $R_\mathrm{s}$, etc.) are estimated perfectly and we use $\alpha = 85\%$ series resistance compensation. For parameters $C_\mathrm{m}^*$ and $R_\mathrm{s}^*$, they are swept with a factor from 0.3 to 3.0; for parameter $C_\mathrm{p}^\dagger$, it is swept from $-4.0\,\mathrm{pF}$ to $4.0\,\mathrm{pF}$; and for parameter $V_\mathrm{off}^\dagger$, it is swept from $-5.0\,\mathrm{mV}$ to $5.0\,\mathrm{mV}$. These values were chosen to show the extreme effects of these imperfect compensations (e.g. we might not expect $C_\mathrm{m}^*$ would be estimated as badly as 3-fold its true value $C_\mathrm{m}$). Although there are no direct measurements for these parameters (these parameters describe machine estimates of true values and error in these), we based this analysis on variability in machine estimates and observed variability in reversal potential in our previous study of the same data Lei et al. (2019).

Figure S3 shows the results of the sensitivity analysis. This simple analysis shows that only parameters $R_\mathrm{s}$ and $V_\mathrm{off}^\dagger$ affect the dynamics of the current, whereas the parameters $C_\mathrm{m}^*$ and $C_\mathrm{p}^\dagger$ affect only a small portion of the recorded current (the transient current that occurs when the voltage is stepped to a new value).

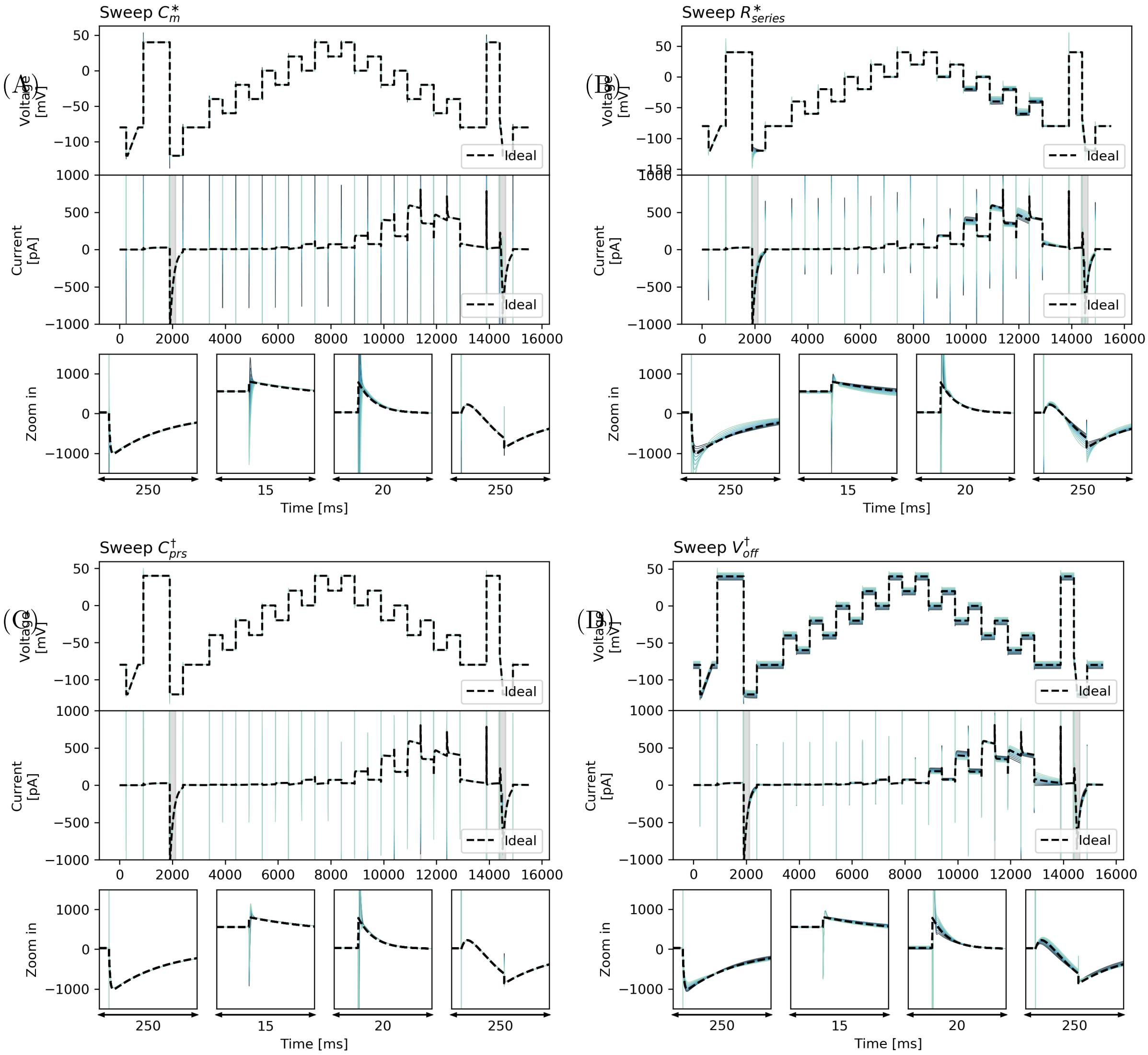


Figure S3: A sensitivity analysis of the voltage-clamp experiment model, showing the effects of imperfect compensation in the voltage-clamp amplifier. Dashed black lines indicate the ideal case, that is $V_{\mathrm{cmd}}$ and $I_{\mathrm{Kr}}$ simulated with $V_{\mathrm{cmd}}$. The voltage traces shown in colour are $V_{\mathrm{m}}$ that the cell sees, and current traces shown in colour are $I_{\mathrm{out}}$ that we observe. Only effects from $V_{\mathrm{off}}$ and $R_{\mathrm{s}}$ are predominant while effects from $C_{\mathrm{m}}$ and $C_{\mathrm{p}}$ have less effect.

### S4 Electrical model cell recordings and simulations

In Figure 4 (main text), only the last 3 s of the staircase protocol is shown. Here in Figure S4, the whole trace recordings and simulation results of the staircase protocol are shown.

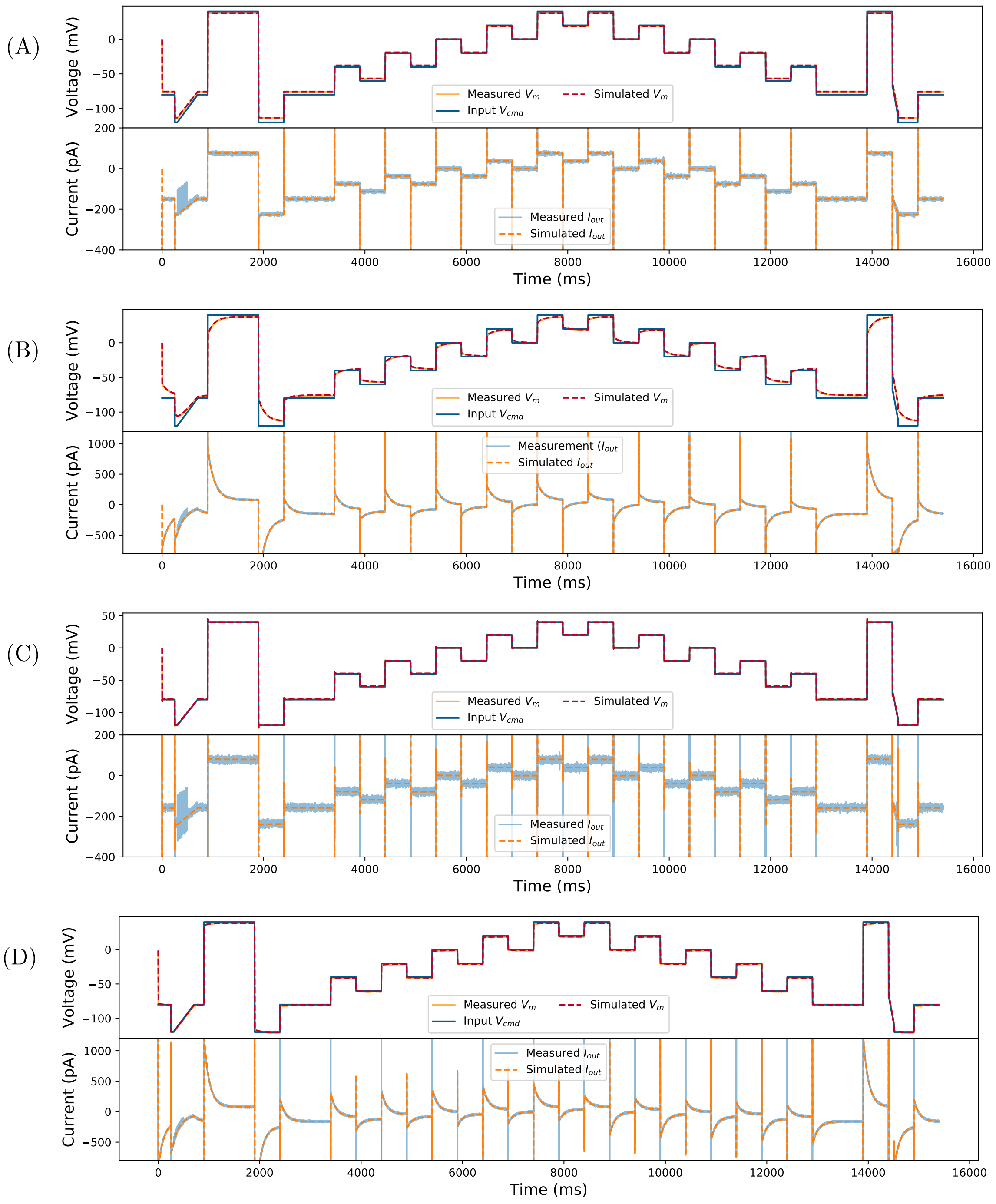


Figure S4: Model simulations (dashed lines) using the amplifier settings compared against the simultaneous voltage clamp-current clamp measurements of the model cells (solid lines). Measurements are shown without compensation using **(A)** Type I Model Cell and **(B)** Type II Model Cell; and measurements with automatic amplifier compensation for $V_{\text{off}}$, $C_{\text{p}}$, $C_{\text{m}}$, and $R_{\text{s}}$ with $\alpha = 80\%$ using **(C)** Type I Model Cell and **(D)** Type II Model Cell. All command voltages were set to be the staircase voltage protocol Lei et al. (2019) (top panel). In the top panel of each subfigure, the blue lines represent the command voltage $V_{\text{cmd}}$, and the orange/red lines represent the membrane voltage $V_{\text{m}}$; the bottom panel shows the current readout via the voltage-clamp, $I_{\text{out}}$.

### S5 Application to Type I Electrical Model Cell

Similar to Section 33.3 in the main text, we use only the uncompensated, raw voltage-clamp measurements (i.e. only $I_{\text{out}}$ and $V_{\text{cmd}}$) to infer the underlying membrane voltage $V_{\text{m}}$ and the parameters of the Type I Model Cell. We then compare the model $V_{\text{m}}$ predictions with the current-clamp measurements. Here, we simply minimise root-mean-squared error (RMSE) using a global optimisation algorithm (CMA-ES) to match the simulated $I_{\text{out}}$ and the recorded current.

Similar to Type II Model Cell results, Figure S5A shows the fitted model $I_{\text{out}}$ (bottom, orange dashed line) and its prediction of the membrane voltage $V_{\text{m}}$ (top, red dashed line), compared with experimental recordings (solid lines). Figure S5B further shows that the fitted model is able to predict current measurements under an independent, unseen voltage-clamp command protocol — a series of action potential waveforms (blue lines in the first panel). Additionally, the agreement between predictions and measurements of $V_{\text{m}}$ (the model is only given the command voltage $V_{\text{cmd}}$) provide assurance the scheme is able to infer the applied $V_{\text{m}}$ as well as predict current.

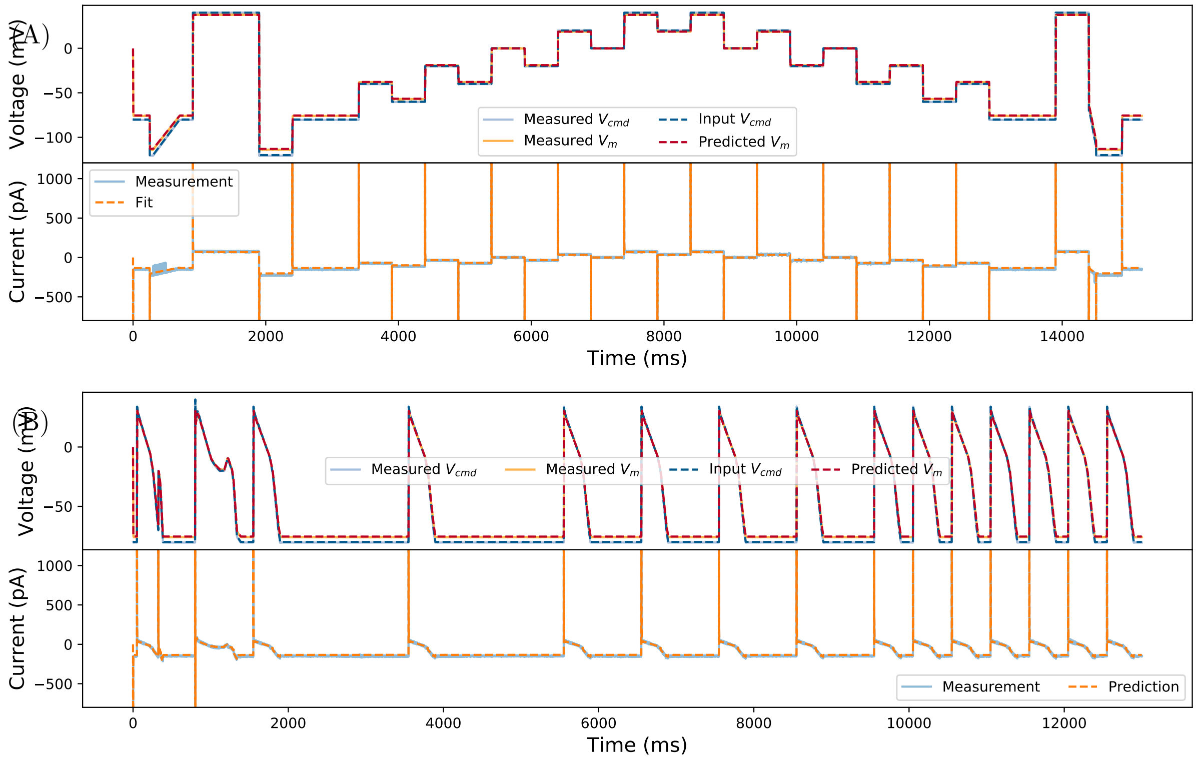


Figure S5: Inferred model simulations and predictions (dashed lines) compared against the experimental data (solid lines) using the Type I Model Cell. **(A)** Model calibration with the staircase protocol (blue lines in the first panel), where model fitted to only the current recording (blue solid line in the second panel). The fitted model was able to predict the membrane voltage $V_{\text{m}}$ (orange solid line), which the model cell sees, measured using the current-clamp. **(B)** Further model validation using an independent voltage-clamp protocol, a series of action potential waveforms (blue lines in the first panel). Again, predictions from the fitted model (dashed lines) closely match both the measured current and membrane voltage.

| | $R_\mathrm{m}$ (MΩ) | $C_\mathrm{p}$ (pF) | $C_\mathrm{m}$ (pF) | $R_\mathrm{s}$ (MΩ) | $V_\mathrm{off}$ (mV) |
|---|---|---|---|---|---|
| Component label | 500.00 | 4.70 | 22.00 | 30.00 | 0.00 |
| Patchmaster estimate | 498.00 | 7.80 | 32.85 | 32.60 | 0.20 |
| Fitted parameters | 567.21 | 23.36 | 32.51 | 34.85 | -0.12 |

Table S1: Type I Model Cell parameters, comparing the values of hardware component labels in the circuit, the values estimated by the Patchmaster amplifier software using a simple test pulse, and our inferred values from the mathematical model.

### S6 Application to simulated patch-clamp data with more complex current kinetics

We then test our parameter inference on the voltage-clamp experiment model with more complex current kinetics. Here we use the $I_{\mathrm{Kr}}$ model described in Lei et al. (2019) (also summarised in Section 44.1 in the main text), to show that with an information-rich protocol, even combined with the voltage-clamp experiment model, theoretically we are able to infer all of the parameters. We first simulated 10 traces of patch-clamp data by using one set of kinetic parameters $\boldsymbol{\theta}$ (identical kinetics assumption, with the hierarchical Bayesian mean parameters from Lei et al. Lei et al. (2019), also shown in Table S4) together with a set of randomly generated voltage-clamp experiment model parameters. The voltage-clamp experiment model parameters were sampled according to

$$V_{\mathrm{off}}^{\dagger} \sim \mathcal{N}(0, 1.5) \qquad (\mathrm{mV}), \tag{S6.21}$$

$$R_{\mathrm{s}} \sim \ln\mathcal{N}(12.5, 2) \qquad (\mathrm{M\Omega}), \tag{S6.22}$$

$$C_{\mathrm{m}} \sim \ln\mathcal{N}(15, 2.5) \qquad (\mathrm{pF}), \tag{S6.23}$$

$$C_{\mathrm{p}} \sim \ln\mathcal{N}(4, 1) \qquad (\mathrm{pF}), \tag{S6.24}$$

$$g_{\mathrm{leak}} \sim \mathcal{N}(0.25, 0.1) \qquad (\mathrm{nS}), \tag{S6.25}$$

$$g_{\mathrm{Kr}} \sim \ln\mathcal{N}(32.3, 32.3) \qquad (\mathrm{nS}). \tag{S6.26}$$

Where $E_{\mathrm{leak}}$ was set to be $-80\,\mathrm{mV}$. We also assumed that the experiments were done with 80% series resistance compensation, and the amplifier estimations have normally distributed errors with a standard deviation of 10%, i.e. after a realisation of $R_{\mathrm{s}}$ we sample $R_{\mathrm{s}}^{*}$ from $\mathcal{N}(R_{\mathrm{s}}, 0.1 * R_{\mathrm{s}})$. Then we perform parameter inference for these synthetic data. Since in actual experiments we know the values of the machine estimations, we do not infer the machine estimates but treat them as known and infer the rest, i.e. $g_{\mathrm{Kr}}, \boldsymbol{\theta}, C_{\mathrm{m}}, R_{\mathrm{s}}, C_{\mathrm{p}}, V_{\mathrm{off}}^{\dagger}$, and $g_{\mathrm{leak}}$.

Figure S6 shows the results of the simulated study with the voltage-clamp experiment model. Figure S6 (Top, Middle) shows the simulated membrane voltage $V_{\mathrm{m}}$ (grey lines) traces differ from the command voltage $V_{\mathrm{cmd}}$ (blue dashed line). The simulated voltage-clamp data are shown in blue with simulated noise, plotted against the fitted model shown in orange. Figure S6 (Bottom) shows the relative error of the inferred parameters. It shows that besides $C_p$, with the fastest dynamics/time scale, all of the parameters can be inferred very accurately even with the simulated noise.

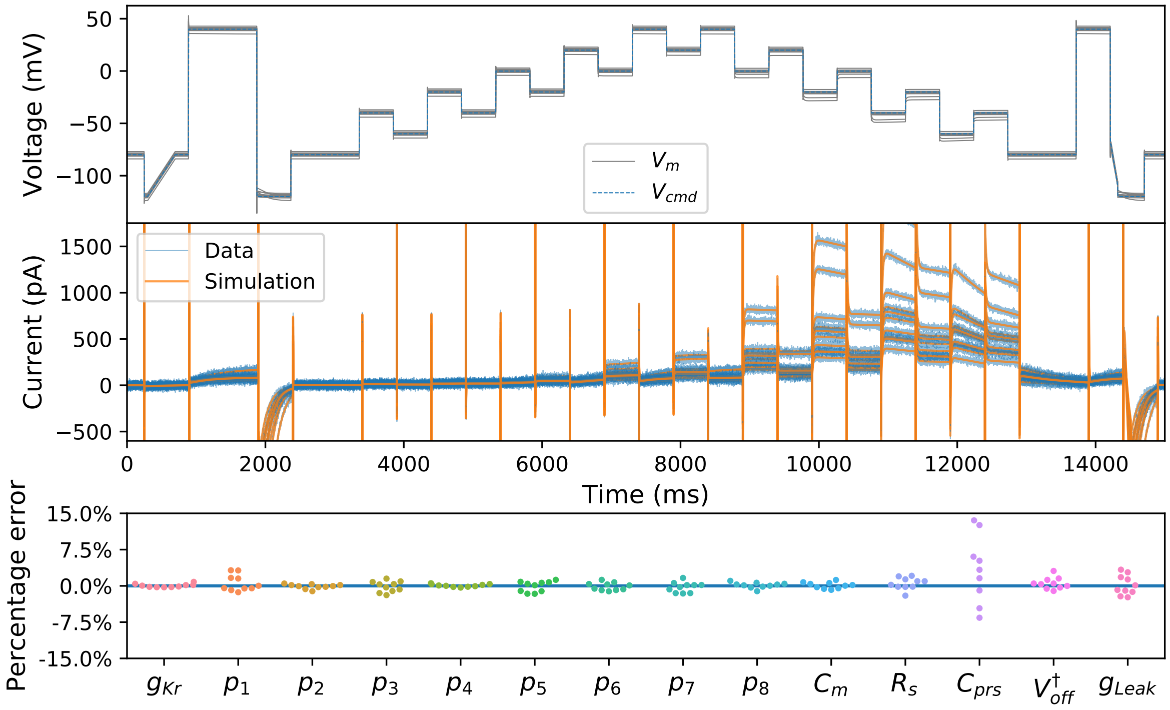


Figure S6: Synthetic $I_{\mathrm{Kr}}$ data study with 10 traces with identical kinetics and different patch artefacts: membrane capacitance, series resistance, seal resistance, pipette capacitance, and leak. The pipette capacitance is the hardest parameter to infer, with errors of up to 15%. But overall, both kinetic parameters and voltage-clamp experiment model parameters are practically identifiable.

### S7 Application to electrical model cell with simplified voltage-clamp experiment model

We perform parameter inference of the simplified voltage-clamp experiment model (Eqs. (18)–(23) in the main text) to the electrical model cell experiments. We focus on the Type II Model Cell as it exhibits nonlinear dynamics and misleads the amplifier into incorrect compensation for $V_{\text{off}}$ (see Table 2 in the main text). Since the simplified voltage-clamp experiment model is based on compensated data, we use amplifier compensated data collected from the Type II Model Cell (Figure S4 (D)).

The most interesting consequence is that due to the dynamical behaviour of this Type II Model Cell, the amplifier estimated both $R_{\text{m}}$ and $V_{\text{off}}$ incorrectly. As shown in see Table 2 in the main text, with a simple test pulse the amplifier estimated $R_{\text{m}}^{*} \approx 90\,\text{M}\Omega$ (when the component $R_{\text{m}} = 500\,\text{M}\Omega$), and $V_{\text{off}}^{*} \approx -1.2\,\text{mV}$ (in the model cell, $V_{\text{off}} = 0\,\text{mV}$). Note that because the amplifier applied (erroneously) $V_{\text{off}}^{\dagger} \approx -1.2\,\text{mV}$, that is the target $V_{\text{off}}^{\dagger}$ for the parameter fit to return.

| | $R_{\text{k}}$ (MΩ) | $C_{\text{k}}$ (pF) | $R_{\text{m}}$ (MΩ) | $V_{\text{off}}^{\dagger}$ (mV) |
|---|---|---|---|---|
| Effective Component Values | 100 | 1000 | 500 | -1.20 |
| Fitted parameters | 96.02 | 1044.68 | 508.34 | -1.40 |

Table S2: Inference of Type II Model Cell parameters with active amplifier compensation using the simplified voltage-clamp experiment model. We compare the components' labelled values in the circuit setup (with $V_{\text{off}}^{\dagger} = -1.2\,\text{mV}$ added alongside as an effective component due to active amplifier compensation), and inferred values from the mathematical model.

Table S2 shows the parameter inference results using the simplified voltage-clamp experiment model. The simplified model is able to: (i) recover Type II Model Cell parameters $R_{\text{k}}$, $C_{\text{k}}$, $R_{\text{m}}$ with good accuracy; and (ii) 're-correct' the amplifier's incorrectly estimated (and thereby artificially introduced) voltage offset. Note that these component estimates are better than the ones from the experiment with no compensation, suggesting a strategy of correcting the amplifier's active compensations may be better than running experiments with no compensation and inferring all these values in postprocessing.

#### S8 Parameter inference algorithm for Hypothesis 2

Optimising $\mathcal{L}$ (Eq. (25) in the main text) is a high-dimensional optimisation problem which can be difficult. To alleviate the bottleneck arising from this problem, we use a parameter inference scheme shown in Algorithm S1, where $L_{\text{kinetics},i}$ is the likelihood of the kinetic parameters $\boldsymbol{\theta}$, and $L_{\text{artefacts},i}$ is the likelihood of $g_{\text{Kr},i}, \boldsymbol{\phi}_i$ for the $i^{\text{th}}$ measurement. By optimising $\mathcal{L}(\boldsymbol{\theta})$, we get $\boldsymbol{\theta}^*, \{g^*_{\text{Kr},i}, \boldsymbol{\phi}^*_i\}_i$, where $\{g^*_{\text{Kr},i}, \boldsymbol{\phi}^*_i\}_i$ vary across measurements which explains all the observed variability.

**Algorithm S1:** Identical kinetic models parameter inference scheme

**Result:** Obtain $\boldsymbol{\theta}^*, \{g^*_{\text{Kr},i}, \boldsymbol{\phi}^*_i\}_i = \text{argmax}_{\boldsymbol{\theta},\{g_{\text{Kr},i},\boldsymbol{\phi}_i\}_i} \mathcal{L}\left(\boldsymbol{\theta}, \{g_{\text{Kr},i}, \boldsymbol{\phi}_i\}_i\right)$

Initialise $\{L_{\text{kinetics},i}\}_i, \{L_{\text{artefacts},i}\}_i$;

Set $\boldsymbol{\theta}^0, \{g^0_{\text{Kr},i}, \boldsymbol{\phi}^0_i\}_i$;

$L^0 \leftarrow \prod_i L_{\text{kinetics},i}\left(\boldsymbol{\theta}^0 | g^0_{\text{Kr},i}, \boldsymbol{\phi}^0_i\right)$;

$j \leftarrow 0$;

**while** *not terminate* **do**

- $\boldsymbol{\theta}^{j+1} \leftarrow$ sample $\boldsymbol{\theta}$ with an optimisation algorithm given $L^j$;
- **foreach** $i$ **do**
  - $g^{j+1}_{\text{Kr},i}, \boldsymbol{\phi}^{j+1}_i \leftarrow \text{argmax}_{g_{\text{Kr},i},\boldsymbol{\phi}_i} L_{\text{artefacts},i}\left(g_{\text{Kr},i}, \boldsymbol{\phi}_i | \boldsymbol{\theta}^{j+1}\right)$
- **end**
- $L^{j+1} \leftarrow \prod_i L_{\text{kinetics},i}\left(\boldsymbol{\theta}^{j+1} | g^{j+1}_{\text{Kr},i}, \boldsymbol{\phi}^{j+1}_i\right)$;
- $j \leftarrow j+1$;

**end**

$\boldsymbol{\theta}^*, \{g^*_{\text{Kr},i}, \boldsymbol{\phi}^*_i\}_i \leftarrow \boldsymbol{\theta}^j, \{g^j_{\text{Kr},i}, \boldsymbol{\phi}^j_i\}_i$

### S9 Remaining relative root mean square error (RRMSE) histograms

Figure S7 shows the relative root mean square error (RRMSE, given by Eq. (26) in the main text) histograms for the remaining validation protocols 1, 2, and 6 that are not included in the main text due to the space limit. Interestingly Hypothesis 1 only shows better predictive performance than Hypothesis 2 for the Validation 2 protocol. We believe this is due to the similarity between Validation 2 and the Calibration protocol — they both step up and down to the same range of voltages in the same duration steps of 500m̀s — i.e. this is the validation situation closest to the training protocol which Hypothesis 1 does better at due to having extra parameters to fit.

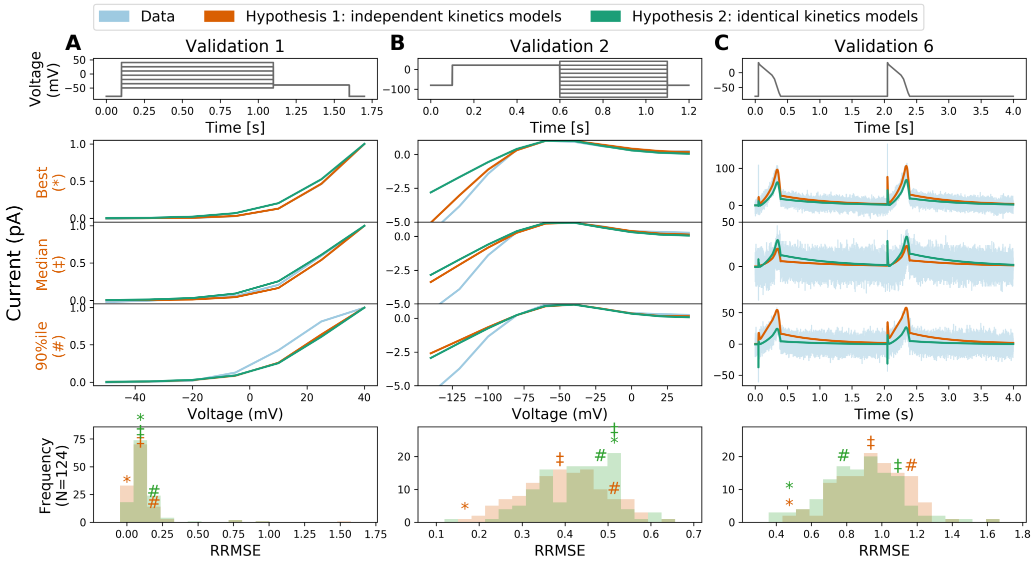


Figure S7: The relative root mean square error (RRMSE, given by Eq. (26) in the main text) histograms for validation protocols 1, 2, and 6, comparing the independent kinetics models from Lei *et al.* Lei et al. (2019) and the identical kinetics models with voltage-clamp artefact. Each histogram represents the same 124 cells with a different protocol and RRMSE each time. Red markers indicate the best (∗), median (‡) and 90$^{\text{th}}$ percentile (#) RRMSE values for the independent kinetics model; green markers are the same cell prediction from the identical kinetics models. For each protocol, the raw traces for the identical kinetics model (green), the independent kinetics model (red), and data (blue) are shown, with the voltage-clamp above. Note that the currents are shown on different scales, to reveal the details of the traces.

### S10 Results for Hypothesis 2: Identical kinetics for all cells with cell-specific artefacts

Table S3 shows the median, $10^{th}$ percentile, and $90^{th}$ percentile of the RRMSE histograms shown in main text Figure 7. Table S4 shows the inferred parameters of the identical kinetics across all 124 cells under the assumptions of Hypothesis 2 (the mean of Hypothesis 1 from Lei et al. Lei et al. (2019) is also shown for comparison). Figure S8 shows the predictions for voltage dependency of steady-states, open probability and time constants using the inferred parameters in Table S4. All of them are calculated directly from inferred parameters using Eqs. (14) & (15) in the main text. Grey lines show the predictions from Hypothesis 1 (assuming cell-specific kinetics with no artefacts; from Lei et al. Lei et al. (2019)), and red lines show the predictions from the mean of Hypothesis 1. Green lines show the predictions from Hypothesis 2 (assuming identical kinetics for all cells with cell-specific artefacts). There is a noticeable shift of the parameters between Hypothesis 1 (mean) and Hypothesis 2. Finally the predictions using the parameters in Table S4 for the staircase protocol is shown in Figure S9, showing the differences in terms of current prediction. The difference is due to the unaccounted experimental artefacts in Hypothesis 1 'contaminating' the physiological (model) parameters, therefore the model parameters from Hypothesis 2 are thought to be more physiologically-relevant.

| | Cal. | | Val. 3 | | Val. 4 | | Val. 5 | | Val. 7 | | Val. 8 | |
|---|---|---|---|---|---|---|---|---|---|---|---|---|
| | H1 | H2 | H1 | H2 | H1 | H2 | H1 | H2 | H1 | H2 | H1 | H2 |
| Median | 0.13 | 0.14 | 0.27 | 0.28 | 1.02 | 0.94 | 0.76 | 0.68 | 0.89 | 0.83 | 0.71 | 0.62 |
| $10^{th}$ %ile | 0.09 | 0.11 | 0.14 | 0.16 | 0.69 | 0.52 | 0.51 | 0.34 | 0.61 | 0.46 | 0.42 | 0.33 |
| $90^{th}$ %ile | 0.17 | 0.20 | 0.52 | 0.65 | 1.43 | 1.38 | 1.26 | 1.17 | 1.26 | 1.31 | 1.11 | 1.05 |

Table S3: The median, $10^{th}$ percentile, and $90^{th}$ percentile of relative root mean square error (RRMSE) histograms shown in Figure 7 in the main text, comparing (**H1**) the independent kinetics models from Lei *et al.* Lei et al. (2019) and (**H2**) the identical kinetics models with voltage-clamp artefact.

| | $p_1$ ($s^{-1}$) | $p_2$ ($V^{-1}$) | $p_3$ ($s^{-1}$) | $p_4$ ($V^{-1}$) | $p_5$ ($s^{-1}$) | $p_6$ ($V^{-1}$) | $p_7$ ($s^{-1}$) | $p_8$ ($V^{-1}$) |
|---|---|---|---|---|---|---|---|---|
| Hypothesis 1 | 7.65e-5 | 9.05e-2 | 2.84e-5 | 4.74e-2 | 1.03e-1 | 2.13e-2 | 8.01e-3 | 2.96e-2 |
| Hypothesis 2 | 1.13e-4 | 7.45e-2 | 3.60e-5 | 4.49e-2 | 9.61e-2 | 2.36e-2 | 7.85e-3 | 3.06e-2 |

Table S4: Inferred parameters for the mean of Hypothesis 1 (assuming cell-specific kinetics with no artefacts; from Lei et al. Lei et al. (2019)) and Hypothesis 2 (assuming identical kinetics for all cells with cell-specific artefacts).

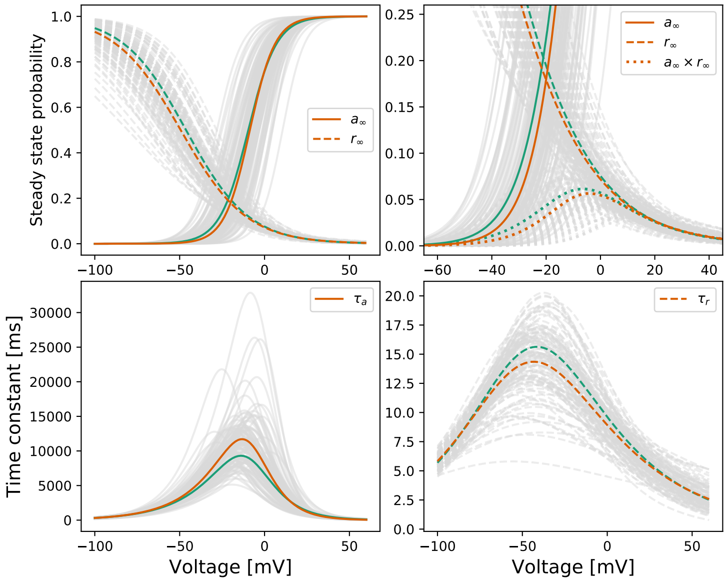


Figure S8: Predictions for voltage dependency of steady-states, open probability and time constants of the model. These lines are calculated directly from inferred parameters using Eqs. (14) & (15) in the main text. Grey lines show the predictions from Hypothesis 1 (assuming cell-specific kinetics with no artefacts; from Lei et al. Lei et al. (2019)), and red/orange lines show the predictions from the mean of Hypothesis 1. Green lines show the predictions from Hypothesis 2 (assuming identical kinetics for all cells with cell-specific artefacts).

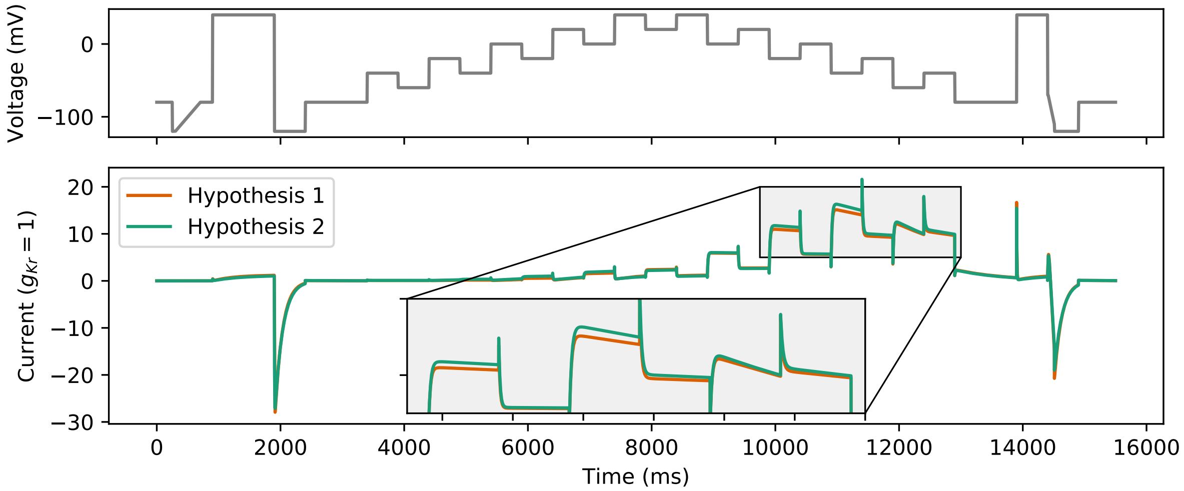


Figure S9: Predictions for the staircase protocol Lei et al. (2019) using the two sets of mean parameters from the two hypotheses in Table S4. The mean of Hypothesis 1, assuming cell-specific kinetics with no artefacts (from Lei et al. Lei et al. (2019)), is shown in red/orange. Hypothesis 2, assuming identical kinetics for all cells with cell-specific artefacts, is shown in green.

### References


Axon Instruments Inc., 1997–1999. Axopatch 200B patch clamp theory and operation. https://www.autom8.com/wp-content/uploads/2016/07/Axopatch-200B.pdf. Accessed: 2020-02-29.

HEKA Elektronik Dr. Schulze GmbH, 2007–2018. EPC 10 USB hardware manual version 2.8. http://www.heka.com/downloads/hardware/manual/m_epc10.pdf. Accessed: 2020-02-29.

Lei, C.L., Clerx, M., Gavaghan, D.J., Polonchuk, L., Mirams, G.R., Wang, K., 2019. Rapid characterisation of hERG channel kinetics I: using an automated high-throughput system. Biophysical Journal 117, 2438–2454.

Moore, J.W., Hines, M., Harris, E.M., 1984. Compensation for resistance in series with excitable membranes. Biophysical Journal 46, 507–14.

Neher, E., 1992. [6] Correction for liquid junction potentials in patch clamp experiments, in: Methods in Enzymology. Academic Press. volume 207 of *Ion Channels*, pp. 123–131.

Neher, E., 1995. Voltage offsets in patch-clamp experiments, in: Sakmann, B., Neher, E. (Eds.), Single-Channel Recording. Springer US, Boston, MA. chapter 6, 2 edition. pp. 147–153.

Sherman, A.J., Shrier, A., Cooper, E., 1999. Series Resistance Compensation for Whole-Cell Patch-Clamp Studies Using a Membrane State Estimator. Biophysical Journal 77, 2590–2601.

Sigworth, F., 1995. Design of the EPC-9, a computer-controlled patch-clamp amplifier. 1. Hardware. Journal of Neuroscience Methods 56, 195–202.

Sigworth, F., Affolter, H., Neher, E., 1995. Design of the EPC-9, a computer-controlled patch-clamp amplifier. 2. Software. Journal of Neuroscience Methods 56, 203–215.

Strickholm, A., 1995. A single electrode voltage, current-and patch-clamp amplifier with complete stable series resistance compensation. Journal of Neuroscience Methods 61, 53–66.

Weerakoon, P., Culurciello, E., Klemic, K.G., Sigworth, F.J., 2009. An Integrated Patch-Clamp Potentiostat With Electrode Compensation. IEEE Transactions on Biomedical Circuits and Systems 3, 117–125.

Weerakoon, P., Culurciello, E., Yang, Y., Santos-Sacchi, J., Kindlmann, P.J., Sigworth, F.J., 2010. Patch-clamp amplifiers on a chip. Journal of Neuroscience Methods 192, 187–92.
